# Spatial transcriptomics reveals organizational properties of the mouse spinal cord and alterations in neuropathic pain

**DOI:** 10.64898/2026.01.10.698734

**Authors:** Qimeng Wang, Yunfei Hu, Yuling Zhu, Gautham Chelliah, Jing Peng, Emmanuella Osei-Asante, Melissa Sobanko, Brianna Sanchez, David D. Ginty, Xin Maizie Zhou, Shan Meltzer

## Abstract

The spinal cord integrates diverse somatosensory inputs and executes motor outputs through anatomically and functionally distinct circuits. Here, we employed spatially resolved single-cell transcriptomic profiling of the adult mouse spinal cord to gain insights into the organizational logic of the spinal cord. Our results reveal distinct spatial and laminar distributions of neuronal subtypes, including axial level and sex-specific differences. Many neuronal subtypes exhibit lamina-specific somatic proximities, suggesting potential regionally specific connectivity. Additionally, neuronal subtypes within the dorsal horn exhibit a wide range of predicted cell-cell communication motifs, as assessed by the spatial distribution of neuropeptide- and other ligand-receptor pairs. Finally, we identified several neuronal subtypes with altered transcriptomic states and predicted cell-cell communications in a model of neuropathic pain. This spatially resolved cellular and molecular map of the spinal cord will facilitate the decoding of circuit mechanisms underlying somatosensory and motor functions in health and disease.--

**Significance statement:** The spinal cord receives and integrates somatosensory inputs and executes motor and autonomic outputs, yet the spatial organization of its diverse cell types in the spinal cord remains incompletely understood. Using spatially resolved single-cell transcriptomics, this study reveals the distribution of spinal cord neurons across axial levels and between sexes, uncovers spatial proximity and predicted communication motifs between neuronal subtypes, and identifies several neuronal populations with transcriptomic changes in a mouse model of neuropathic pain. These findings provide a spatial framework for linking molecularly defined spinal cell types to circuit function and pain. The resource generated in this study will guide future molecular and functional studies of spinal cord somatosensory, motor, and autonomic circuits in health and disease.

## Introduction

The spinal cord is a critical gateway for information transfer between the brain and body, as well as a processing center with neuronal subtypes that integrate somatosensory information and execute motor and autonomic functions (1, 2). These processing and transfer functions rely on a wide array of spinal cord cell types with distinct laminar distributions, molecular signatures, morphologies, neurotransmitters, physiological properties, and, in some cases, vulnerabilities to pathological conditions (3, 4). The basic anatomical unit of the spinal cord is a segment, within which sensory afferents convey information from the periphery, autonomic and motor efferents control muscles and other organ systems, and ascending projection neurons relay signals to the brain. Along the rostrocaudal (R-C) axis, sensory inputs and motor pool function are distributed over multiple segments. Along the dorsoventral (D-V) axis, the primary functional organization of the mammalian spinal cord exhibits regional specification: the dorsal horn processes sensory inputs and relays their signals to the brain and intermediate and ventral spinal cord; the intermediate zone integrates and processes sensorimotor signals and also coordinates ascending and autonomic outputs; and the ventral horn executes motor functions. Local spinal processing is shaped by descending inputs from supraspinal centers, which modulate both sensory processing and control motor outputs (5, 6). However, the organizational logic of interneurons, projection neurons, and motoneurons across the different axial levels of the spinal cord, which interact with distinct dermatomes, internal organs, and muscle groups, remains incompletely understood.

Primary somatosensory neurons convey a rich repertoire of signals encoding tactile, nociceptive, thermal, chemical, and proprioceptive stimuli via their central axons that enter the spinal cord dorsal horn and synapse onto select spinal interneurons, projection neurons, and motoneurons (7, 8). Nociceptor and thermosensitive afferents predominantly terminate within dorsal horn superficial laminae I-II, innocuous mechanosensitive afferents project to laminae II_iv_-V, and proprioceptive afferents target the intermediate zone and ventral horn, where pre-motor interneurons and motoneurons (MNs) are situated. Sensory inputs from the internal organs, such as the stomach, colon, bladder, and genitalia, exhibit axial-level selectivity (9, 10). Moreover, interneurons throughout the spinal cord integrate peripheral sensory inputs, thereby shaping ascending signals, as well as motor and autonomic outputs (11). As such, the precise positioning of spinal cord neurons across dorsal-ventral, medial-lateral, and axial levels of the spinal cord dictates the specificity with which somatosensory inputs can form to engage local circuits critical for somatosensation as well as autonomic and motor control.

Several recent single-cell (scRNA-seq) and single-nucleus RNA sequencing (snRNA-seq) studies have revealed an impressive molecular and cellular heterogeneity of the spinal cord, uncovering a myriad of cell types and their marker genes (12–18). However, the distribution of cell types across cervical, thoracic, lumbar, sacral, and coccygeal levels has remained largely unexplored (1). Furthermore, although males and females differ in somatosensory processing (19), and targeted studies of selected cell populations suggest underlying sex differences within the spinal cord (20), the identities and axial-level distributions of sexually dimorphic cell types remain poorly understood.

Dysfunction of the spinal cord is implicated in a wide array of neurological disorders that span sensory, motor, and autonomic domains (21, 22). Neuropathic pain, for example, which afflicts 3%–17% of the general population, is marked by allodynia—pain evoked by normally innocuous stimuli such as gentle touch (23–25). Following peripheral nerve injury, the spinal cord undergoes reorganization at molecular, cellular, and circuit levels, which underlies the generation and “chronification” of central sensitization leading to allodynia (4, 26, 27). Current treatments for neuropathic pain have moderate efficacy, and new therapeutic targets and approaches are needed (23). A better understanding of the spatially resolved molecular and cellular changes in the spinal cord will open new avenues for treating neuropathic pain and other disorders of spinal cord function.

Here, we used multiplexed error-robust fluorescence in situ hybridization (MERFISH), an image-based method for spatially resolved single-cell transcriptomics (28), to investigate the spatial organization of cell types and their gene expression in the spinal cord of healthy mice across axial levels and between sexes, as well as in a neuropathic pain state. We uncovered candidate neuropeptide and other ligand-receptor interactions with their cognate receptors across spinal cord cell subtypes. Furthermore, neuropathic pain was associated with molecular and cellular changes, including the level of activity-regulated genes and predicted neuropeptide interactions, in several dorsal horn neuronal subtypes. Together, this study provides a rich resource for evaluating spinal cord cell types and circuits that underlie somatosensation, movement, and autonomic control across axial levels and between sexes in healthy animals and in a neuropathic pain state.

## Results

### Diversity and organization of cell types in the adult mouse spinal cord

To elucidate the cellular composition and spatial organization of the adult mouse spinal cord, we employed MERFISH, an imaging-based approach for spatially resolved single-cell transcriptomics with error-robust barcoding reads (28). MERFISH detects the location of individual RNA molecules within tissues, which can reveal the spatial organization of cell types across anatomically defined regions (29–31). Based on previous spinal cord scRNA-seq and snRNA-seq studies (12–14), we designed a panel of 500 genes that includes combinatorial markers for spinal cord cell types, functionally relevant genes encoding neurotransmitters, neuropeptides, ligand-receptor pairs, and neuronal activity-regulated genes (Table S1; see details in Materials and Methods). We collected and imaged transverse spinal cord sections spanning cervical, thoracic, lumbar, and sacral levels from five male and five female mice (C57BL/6J, 8-10 weeks old). The imaged RNA species were decoded, spatially mapped, and assigned to segmented cells (Figure S1A). Overall, we obtained 146,621 high-quality cells in spinal cords from six independent replicates with high consistency across replicates (Figure S1B). Furthermore, expression of individual genes correlated well with bulk RNA-seq results from spinal cords (Figure S1C) (32). Spatial distributions of marker genes, such as *Chat*, *Pdyn*, and *Tac2*, are detected in regions consistent with in situ hybridization (ISH) data from the Allen Spinal Cord Atlas (Figure S1D).

Unsupervised clustering revealed 14 major cell types: five neuronal classes, including dorsal excitatory neurons, dorsal inhibitory neurons, medial-ventral excitatory neurons, medial-ventral inhibitory neurons, cholinergic neurons, and nine non-neuronal classes, which include oligodendrocytes, oligodendrocyte precursor cells (OPCs), oligodendrocyte progenitors, astrocytes, microglia, endothelial cells, meningeal cells, ependymal cells, and peripheral glial cells (Figure 1A). These major cell types were further subdivided into a total of 60 transcriptionally distinct subtypes: 24 excitatory neuron subtypes, 15 inhibitory neuron subtypes, 3 skeletal motoneuron subtypes, visceral motoneurons (preganglionic autonomic neurons), cholinergic interneurons, and 16 non-neuronal subtypes (Figures 1B and 1C; see details in Materials and Methods). We named the neuronal subtypes using a three-element nomenclature. The first element denotes their general anatomical location (i.e., DH for dorsal horn, DM for dorsal to intermediate, M for intermediate, and MV for intermediate to ventral). The second element denotes whether they are excitatory (ex) or inhibitory (in), and the third element is their family and/or subcluster marker gene. Among non-neuronal classes, oligodendrocytes, astrocytes, and oligodendrocyte progenitors each exhibited two subtypes, and meningeal cells had three subtypes (Figure 1C) (12, 13, 33).

**Figure 1.**
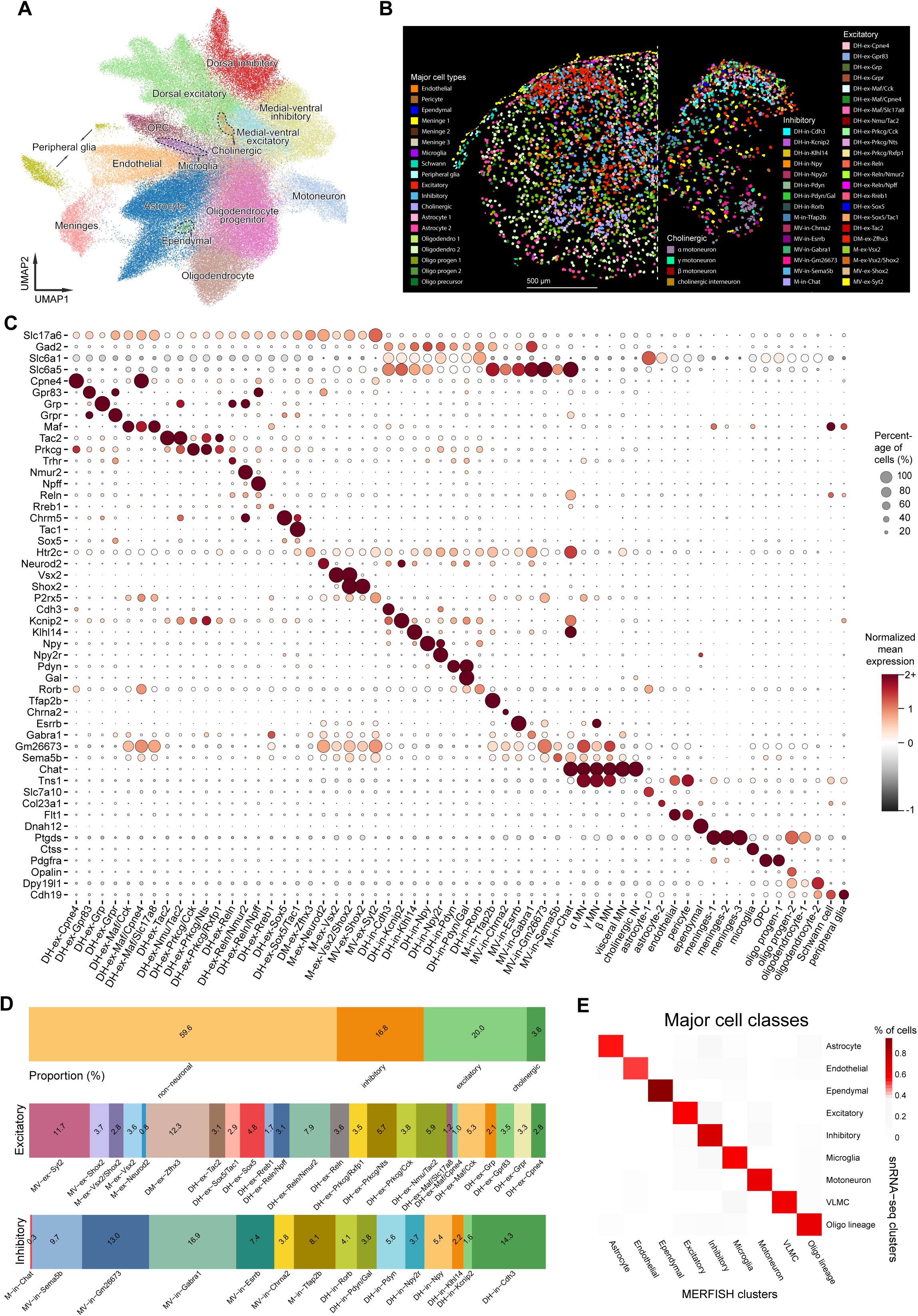
MERFISH reveals molecularly diverse cell types and subtypes comprising the mouse spinal cord. (A) UMAP visualization of all cells (number = 146,621, 5 males and 5 females) identified by MERFISH, color-coded by major cell classes. (B) Spatial distribution of MERFISH-determined cell types is shown on a representative transverse section of the cervical spinal cord. Scale bar, 500 μm. (C) Dot plot of gene expression of all molecularly defined cell types. Color indicates the level of normalized mean expression, and dot size represents the percentage of cells expressing each gene. (D) Stacked bar charts of proportions of cell categories (top), excitatory neurons (middle), and inhibitory neurons (bottom). (E) Heatmap of the percentage of MERFISH major cell classes assigned to each single-nucleus RNA sequencing (snRNA-seq) cell cluster label.

Using the cell coordinates, we projected these clusters in space, which revealed the anatomical layout of the transverse section and the precise location of each cell (Figure 1B). Among excitatory and inhibitory neurons, dorsal horn subtypes exhibited notable laminar distribution patterns (Figures S2A and S2B). Both excitatory and inhibitory neuron subtypes in the dorsal horn were confined to defined laminae (Figures S2A and S2B). For example, the DH-ex-Gpr83 subtype, which we speculate to consist of projection neurons, was largely confined to the superficial lamina (34), and the DH-in-Cdh3 subtype was localized to lamina III (7). Neuronal subtypes in the intermediate and ventral regions of the cord displayed more diffuse distributions without clear laminar boundaries (Figures S2A and S2B). As expected, cholinergic interneurons and motoneurons were localized in the intermediate zone and the ventral horn, respectively (Figure S2C). Most non-neuronal cells were broadly distributed throughout the spinal cord, except for ependymal cells, which surrounded the central canal, one type of oligodendrocyte and one type of astrocyte, both found in white matter tracts, and peripheral glia, Schwann cells, and meningeal cells (Figures 1B and S2D).

The second round of unsupervised clustering revealed known families of neuronal subtypes (Figure 1C) (12–18). The Prkcg excitatory neuron family, comprising DH-ex-Prkcg/Nts, DH-ex-Prkcg/Cck, and DH-ex-Prkcg/Rxfp1 subtypes, contains neurons mediating light touch and allodynia under pathological conditions (35). Subtypes of the Maf excitatory neuron family, DH-ex-Maf/Cck, DH-ex-Maf/Slc17a8, and DH-ex-Maf/Cpne4, are associated with light touch mechanoreception (36–38). The Reln excitatory neuron family, including DH-ex-Reln, DH-ex-Reln/Npff, and DH-ex-Reln/Nmur2, mediates transmission of chemical itch (13). DH-in-Pdyn and DH-in-Pdyn/Gal subtypes express the marker gene encoding prodynorphin, which is also involved in mechanical pain and chemical itch transmission (39, 40). Neuronal subtypes of the Sox5 family, including DH-ex-Sox5 and DH-ex-Sox5/Tac1, are likely involved in mechanical nociception (13). Several additional neuronal subtypes of the dorsal horn with known anatomical features and physiological functions also emerged. For example, DH-ex-Grpr neurons, corresponding to Grpr^+^ vertical cells implicated in both itch and pain, reside in the superficial lamina (41). DH-ex-Gpr83 are likely anterolateral pathway projection neurons that convey tactile information to higher brain centers (34). Inhibitory neuron subtypes primarily in lamina III, including the DH-in-Rorb, DH-in-Cdh3, DH-in-Npy, and DH-in-Kcnip2 subtypes, exhibit unique morphological and physiological properties and contribute to light touch information processing (42–44). Among neuronal subtypes in the intermediate and ventral zones, MV-in-Chrna2 neurons, marked by expression of *Chrna2*, *Chrna7*, and *Calb1*, correspond to Renshaw cells, a class of inhibitory interneurons derived from the V1 lineage mediating recurrent inhibition of motoneurons (45). MV-in-Gm26673 neurons likely correspond to Ia inhibitory neurons (13). MV-ex-Shox2 neurons participate in rhythm generation for locomotion as part of the central pattern generator (46). M-ex-Vsx2 and M-ex-Vsx2/Shox2 neurons are likely derived from the V2a lineage and involved in skilled motor programs controlling reaching (47).

Unsupervised clustering of the cholinergic neuron populations revealed skeletal motoneurons, visceral motoneurons, and cholinergic interneurons (Figures S2E-S2K). Skeletal motoneurons, residing in the spinal cord ventral horn with high expression of *Chat* and *Tns1* (Figure S2F) (33), were subdivided into three neuronal subclusters, including α (marked by *Rbfox3* and *Vipr2*; Figure S2G), γ (marked by *Esrrg* and *Esrrb*; Figure S2H), and a subtype without a known marker, which we speculate are the elusive β motoneurons (48). Visceral motoneurons (preganglionic autonomic neurons; Figure S2I), marked by *Nos1*, were located in the lateral horn, and a group of cholinergic interneurons, marked by *Pitx2* and *Pax2* (Figure S2J), were located near the central canal (Figure S2K) (33, 49). Cerebrospinal fluid-contacting neurons were not detected, likely due to the low detection of the *Pkd2l1* marker gene (50). Overall, non-neuronal cells were most abundant and accounted for 59.6% of the total spinal cord cell population, followed by glutamatergic excitatory neurons (20.0%), inhibitory neurons (16.8%), and cholinergic neurons (3.6%; Figure 1D). Proportions of major cell classes and neuronal subtypes were consistent across 10 biological replicates (Figures S3A-S3C). Together, the spatial and transcriptional information in the MERFISH dataset allowed us to connect spatially-defined neuronal subclusters to unique anatomical and functional neuronal subtypes in the spinal cord.

To assess the detection accuracy of MERFISH profiling, we performed an integrated analysis of our MERFISH data with mouse spinal cord snRNA-seq data (14). All major cell types exhibited a strong correlation between the two datasets (Figures 1E, S3D, and S4A-S4C). MERFISH recovered more cholinergic neurons compared to previous snRNA-seq studies (Figure S3D). The neuronal subtypes and their spatial distribution in our MERFISH data also correlated well with the neuronal groups reported in a recent Xenium dataset of the mouse spinal cord (51) (Figure S4D and S4E). Consistent with the use of MERFISH in other tissues (52, 53), our analysis classified many of the snRNA-seq-defined clusters at finer resolution with more subclusters, possibly because of their higher rates of detection using MERFISH (Figure S4C).

### Heterogeneous distribution of neuron subtypes along the rostral-caudal and dorsal-ventral axes of the spinal cord

The spinal cord exhibits regional specialization along both the rostral-caudal and dorsal-ventral axes with distinct functional domains corresponding to different body regions (R-C axis), and sensory and motor circuits (D-V axis) (3, 54). To discern the spatial organization of neuronal subtypes along the R-C axis, we investigated the relative abundance of subtypes in each axial level (Figure 2A). Heterogeneous distributions of neuronal subtypes along the R-C and D-V axes were evident. Along the R-C axis, a relative enrichment of many neuronal subtypes was observed at all axial levels. As expected, our analysis showed that visceral motoneurons were concentrated at thoracic and lumbar levels (Figure 2B), where they formed the intermediolateral column comprising the preganglionic sympathetic autonomic output pathway (55). The analysis also revealed previously unreported regional enrichments, such as selective enrichment of M-ex-Neurod2 neurons at thoracic and lumbar levels and M-ex-Vsx2 neurons at lumbar and sacral levels (Figure 2B). Intriguingly, 9 of the 13 neuronal subtypes enriched at the thoracic level were excitatory neurons, while 8 out of the 11 subtypes enriched at the sacral level were inhibitory neurons (Figure 2B, Table S2).

**Figure 2.**
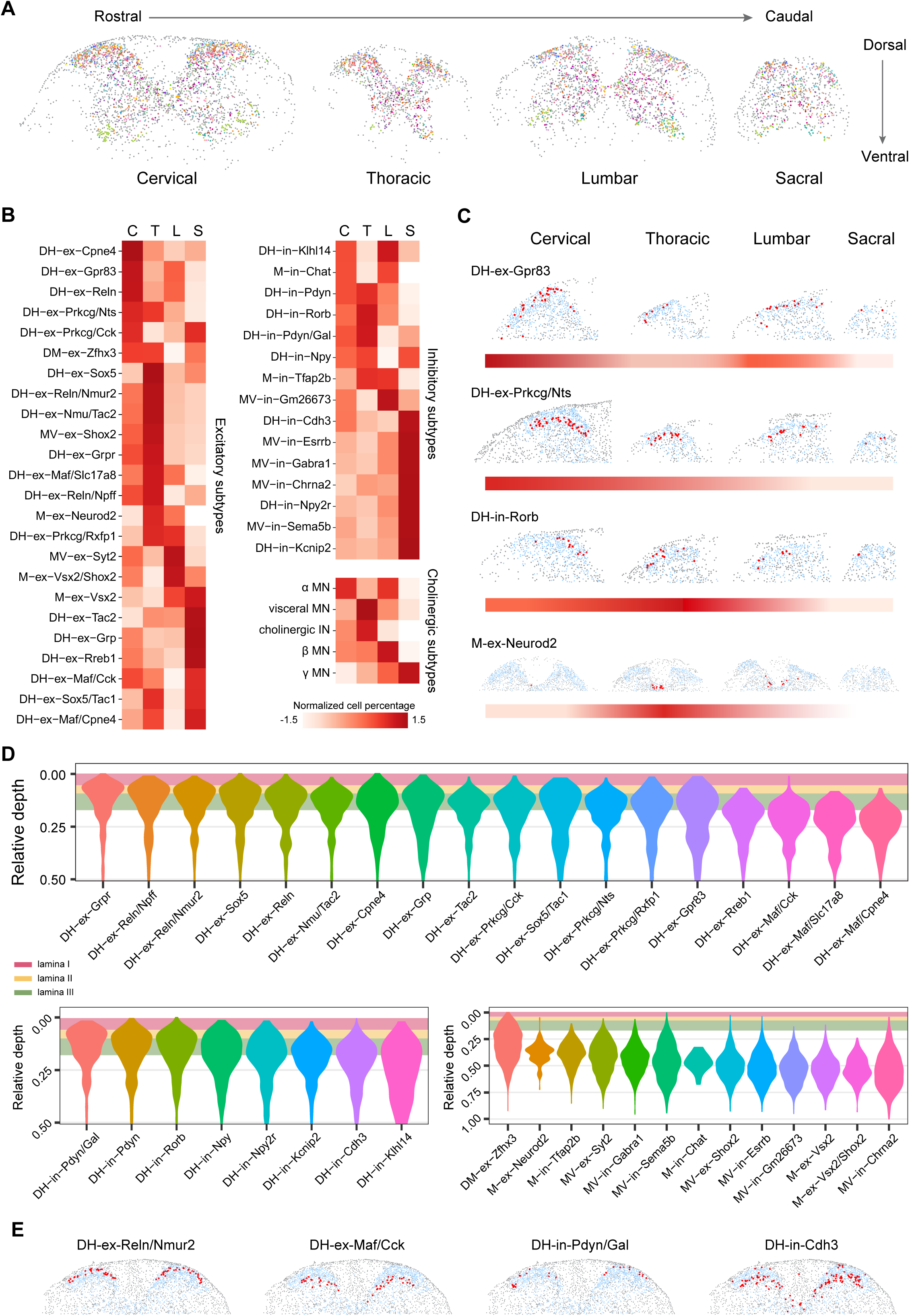
Spatial organization of different neuronal subtypes in the spinal cord. (A) Serial transverse MERFISH sections showing the spatial organization of neuron subtypes along rostral-caudal (R-C) and dorsal-ventral (D-V) axes. (B) Heatmaps showing normalized proportions of neuron subtypes at cervical (C), thoracic (T), lumbar (L), and sacral (S) levels. The normalized cell percentage is calculated using a z-score. (C) Spatial distributions of example neuronal subtypes on representative sections at different axial levels. Red dots mark the indicated subtype of neurons, light blue dots mark the rest of the neurons, and gray dots mark non-neuronal cells. The color gradient indicates the relative proportion of the respective neuron subtype at the respective axial level. (D) Violin plots showing distributions of the relative depth of respective neuron subtypes across all axial levels, calculated as the ratio of the distance of neurons to the dorsal surface of the spinal cord to the maximum distance of the section. (E) Spatial locations of DH-ex-Reln/Nmur2, DH-ex-Maf/Cck, DH-in-Pdyn/Gal, and DH-in-Cdh3 neuron subtypes on the representative cervical section.

Among excitatory neuron populations, most subtypes were regionally enriched in a modular manner rather than following gradual transitions between regions, with a few exceptions. DH-ex-Tac2 neurons, for instance, were gradually enriched from rostral to caudal levels, whereas enrichment of DH-ex-Prkcg/Nts neurons followed the reverse order (Figure 2B). In contrast, nearly half of the inhibitory neuron subtypes peaked in abundance at the sacral level (Figure 2B). Most cholinergic neuron subtypes were less abundant at the sacral level, while α motoneurons were enriched at both cervical and lumbar levels (Figure 2B), which is consistent with their selective innervation of extrafusal fibers of muscle spindles (56). It is noteworthy that three neuronal subtypes, M-ex-Neurod2, M-in-Chat, and cholinergic interneurons, were absent at the sacral level. Spatial projections of various neuronal subtypes on transverse spinal cord sections showed their uneven distributions along the R-C axis (Figure 2C).

Another readily recognizable feature was the laminar organization of neuronal subtypes along the D-V axis, especially within the dorsal horn (Figure 2A). Calculation of the relative depth from the dorsal surface of the spinal cord section revealed the laminar distribution of individual neuronal subtypes. To delineate the ventral boundaries of superficial laminae, we also collected and stained sections adjacent to MERFISH sections for CGRP and Kv4.3 and calculated their relative depths (Figure 2D). Neuronal subtypes in the dorsal horn were more confined within laminae, whereas subtypes in intermediate and ventral regions were more diffusely distributed. For example, the *Reln* populations were primarily present superficially within lamina I. *Prkcg* family neurons (i.e., DH-ex-Prkcg/Cck, DH-ex-Prkcg/Nts, and DH-ex-Prkcg/Rxfp1 subtypes) were located in laminae II and III, and the *Maf* populations (i.e., DH-ex-Maf/Cck, DH-ex-Maf/Slc17a8, and DH-ex-Maf/Cpne4 subtypes) were located in laminae III-IV. Similarly, the dorsal inhibitory DH-in-Pdyn/Gal subtype resided predominantly in lamina I, and DH-in-Pdyn and DH-in-Rorb subtypes resided primarily across laminae I and II and occasionally in lamina III (39, 40, 44). DH-in-Cdh3 and DH-in-Npy subtypes largely resided deeper in lamina III (7). Plotting each subtype onto representative cervical sections revealed their highly specific spatial localizations in the spinal cord across all axial levels (Figures 2E and S2A-S2C). Plotting each subtype at individual axial levels revealed several neuron types with broader or more restricted distribution in the spinal cord. For example, M-in-Tfap2b and MV-ex-Shox2 subtypes exhibited more restricted distributions at thoracic and lumbar levels (Figure S4F).

Most non-neuronal cell types exhibited a broad distribution throughout the spinal cord with a few exceptions (Figure S2D). Astrocyte-1 cells, marked by the expression of *Slc7a10*, were primarily found in the grey matter, while astrocyte-2 cells, marked by the expression *Col23a1*, and oligodendrocyte-2 cells, marked by *Cyp27a1* and *Rab37*, were present only in the white matter (Figure S2D).

Together, the spatial analyses revealed that neuronal subtypes in the spinal cord were organized in highly structured yet regionally heterogeneous patterns along both R-C and D-V axes, reflecting distinct molecular and functional subdomains across axial levels and laminae.

### Distinct spinal neuron subtypes are differentially enriched in males and females

The spinal cord exhibits sex differences in nociceptive information processing, tactile reactivity, and motor control circuits (55, 57, 58), suggesting that sexual dimorphism may contribute to segment-specific function and disease susceptibility. However, sex-specific differences in spinal cord neuron subtypes across axial levels remain poorly understood. To this end, we compared the relative abundance of each neuronal subtype between males and females across cervical, thoracic, lumbar, and sacral levels (Figure 3A). 11 of the 44 neuronal subtypes exhibited significant differences in the percentages of cells across axial levels between males and females. Among the five dorsal horn neurons showing sex differences, DH-in-Npy2r neurons were more enriched in cervical segments in females (Figure 3B), whereas DH-ex-Reln/Nmur2 neurons were more abundant in lumbar segments in males (Figure 3C). M-ex-Neurod2 neurons exhibited an enrichment at the thoracic level in males (Figure 3D), while MV-in-Gm26673 neurons were more abundant in lumbar sections in females (Figure 3E). Intriguingly, visceral motoneurons were enriched in males at the lumbar level (Figure 3F), and γ motoneurons showed a female-biased increase in lumbar regions (Figure 3G), indicating sex-specific physiological controls of internal organ function and motor outputs.

**Figure 3.**
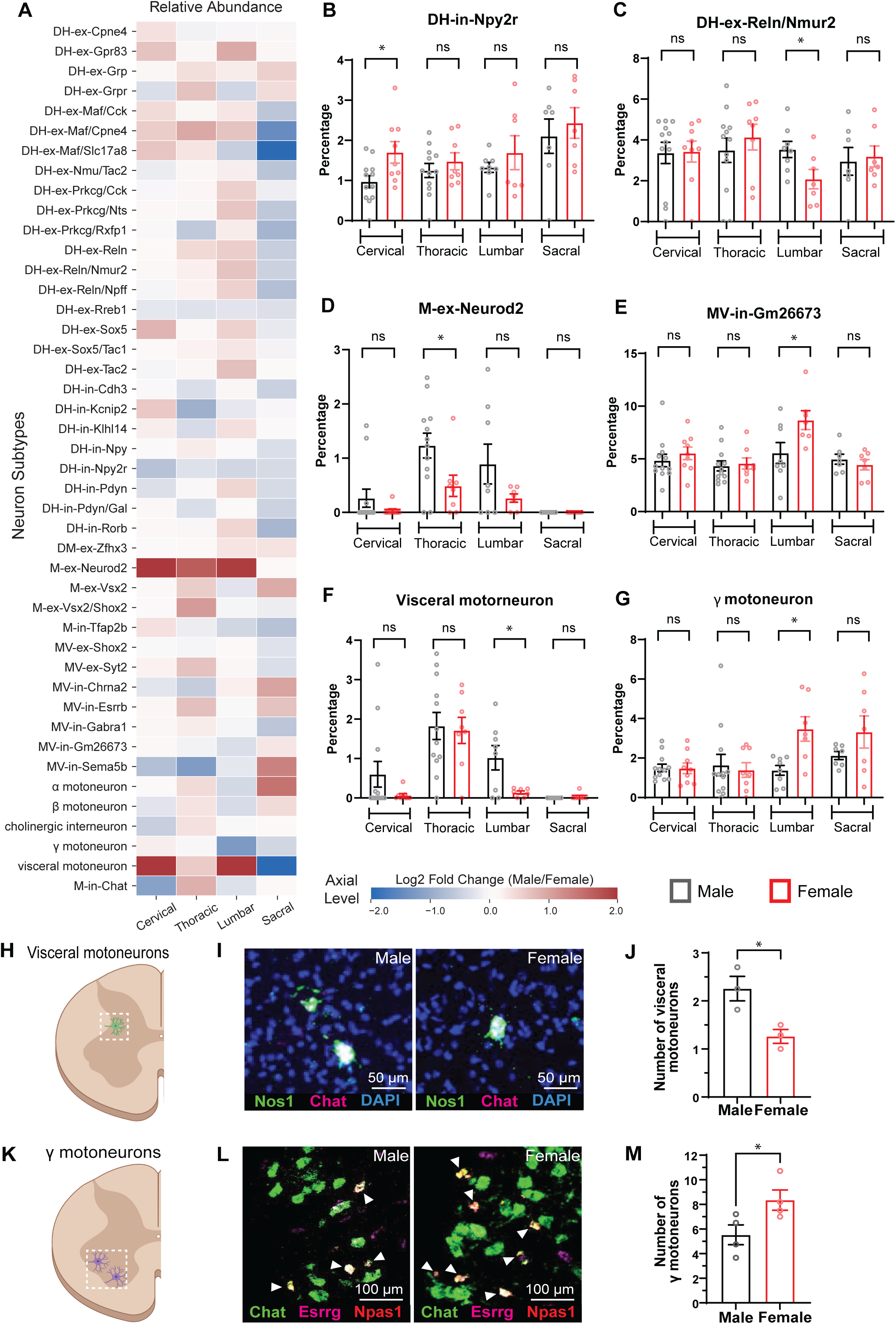
Sex differences in the proportion of neuronal subtypes across spinal cord axial levels. (A) Heatmaps showing the relative abundance of neuron subtypes at cervical (C), thoracic (T), lumbar (L), and sacral (S) levels in males and females. (B-G) Quantification of DH-in-Npy2r (B), DH-ex-Reln/Nmur2 (C), M-ex-Neurod2 (D), MV-in-Gm26673 (E), visceral motoneuron (F), and γ motoneuron (G) subtypes as a percentage of total neurons per section. Welch’s t-tests. (H) Schematic of the location of visceral motoneurons in the spinal cord. (I) Fluorescent *in situ* hybridization images of visceral motoneurons as co-labeled with *Nos1* and *Chat*. (J) Quantification of *Nos1* and *Chat* double-positive neurons per section in wildtype animals (32 sections from 3 males and 26 sections from 3 females). (K) Schematic of the location of γ motoneurons in the lumbar spinal cord. (L) Fluorescent *in situ* hybridization images of γ motoneurons as co-labeled with *Chat, Esrrg, and Npas1*. (M) Quantification γ motoneurons in wildtype animals (29 sections from 4 males and 28 sections from 4 females). Each dot represents a data point from one animal. Student’s t-test; ns, not significant; ∗p < 0.05; error bars indicate standard error of the mean (SEM).

To validate the observed male-biased enrichment of visceral motoneurons in the lumbar spinal cord, we performed RNAscope *in situ* hybridization using probes against *Chat* and *Nos1* to locate visceral motoneurons and quantified their numbers within the intermediolateral autonomic nucleus (Figures 3H and 3I). We also visualized γ motoneurons in the lumbar spinal cords using *Chat*, *Esrrg*, and *Npas1* as markers (33) and quantified the number of γ motoneurons within the ventral horn in both males and females (Figures 3K and 3L). Consistent with the MERFISH analysis, we observed a significant increase in the number of visceral motoneurons and a significant decrease in the number of γ motoneurons in males compared to females (Figures 3J and 3M). Together, these findings suggest that sexually dimorphic organization of spinal circuits may underlie sex differences in sensory, autonomic, and motor functions.

### Cell-cell proximity analysis predicts lamina-specific somatic interactions in a cell-type-specific manner

The spinal cord integrates multi-modal sensory inputs in local circuits for somatosensory information processing (59, 60). High-resolution MERFISH measurements of the spatial locations of the cells have enabled the prediction of cell-cell interactions arising from somatic contact, which can be inferred when cell pairs occur in close spatial proximity more often than expected by random distribution (53, 61). Therefore, we asked whether our MERFISH dataset may reveal cell type-specific cell-cell proximity in the spinal cord and potential interaction pairs. We analyzed the cell type composition within the local “neighborhood” of each cell in the spinal cord and calculated the extent of cell type colocalization in space (Figure 4A). Many pairs of cells displayed enrichment or a lack of spatial proximity. As expected, motoneuron subtypes were in close proximity to each other in the ventral horn. Most glial cell types, on the other hand, did not exhibit proximity to one another, consistent with their tiled organization (Figure 4A) (62).

**Figure 4.**
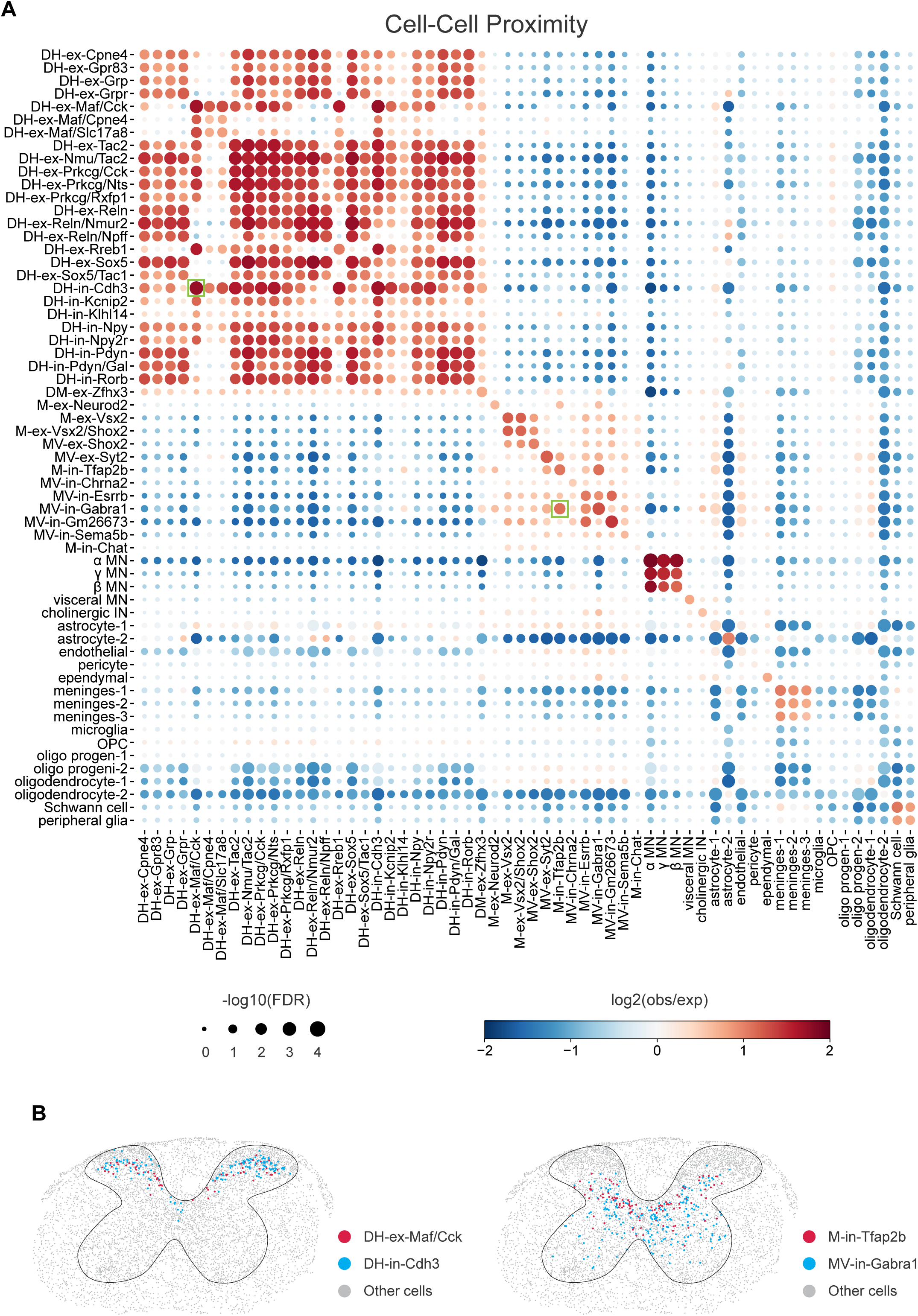
Cell-cell proximity across spinal cord cell subtypes. (A) Enrichment of spatial proximity between pairs of cell subtypes shown in a dot plot. Proximity is computed from soma centroid locations. Color represents the log -transformed ratio of observation to expectation of colocalization frequency, indicating enriched (red) or depleted (blue) proximity of the two cell subtypes. Dot size represents the –log -transformed false discovery rate (FDR)-adjusted *p*-value, reflecting the statistical significance of colocalization. Green boxes denote the example pairs shown in panel (B). (B) Spatial distributions of DH-ex-Maf/Cck and DH-in-Cdh3 subtypes (left), and M-in-Tfap2b and MV-in-Gabra1 subtypes (right) on representative sections. The black line delineates the grey matter.

Many neuronal subtypes in the dorsal horn exhibited close proximity in space in a lamina-specific manner, which may be predictive of local neural microcircuits, such as excitatory–inhibitory coupling. For example, projection neurons DH-ex-Gpr83 exhibited high proximity to DH-ex-Reln/Nmur2 and DH-ex-Sox5 neurons. DH-ex-Grpr vertical cells in lamina I/II were in close proximity to DH-ex-Reln/Nmur2 and DH-ex-Nmu/Tac2 neurons in lamina II. Within lamina III, DH-in-Cdh3 neurons exhibited strong spatial proximity to DH-ex-Maf/Cck neurons (Figure 4B), suggesting potentially higher connectivity between these two interneuron subtypes in modulating tactile information processing (42). Furthermore, interneurons of the *Maf* populations in lamina III showed selective proximity to only a few other cell types, such as DH-ex-Rreb1 neurons. In the intermediate zone, a few pairs of neuronal subtypes showed elevated spatial proximity, including the M-in-Tfap2b and MV-in-Gabra1 subtypes (Figure 4B). Intriguingly, astrocyte-1 cells in the gray matter showed higher proximity to MV-in-Gabra1 neurons in laminae V-VII (Figure 4A). Thus, MERFISH facilitated the recognition of spatial proximity with cell-type specificity, which can be further investigated using histological and physiological approaches to determine the functional significance of the spatial proximity between selected neuronal subtypes in the spinal cord.

### Distinct cell-cell communication motifs within and between dorsal horn laminae of the spinal cord

How neurons send, receive, and respond to chemical signals from neighboring cells, through cell-cell communication (CCC), is fundamental to information transfer within the nervous system. Classic neurotransmitters, neuropeptides, and a range of other ligands, including neurotrophins and semaphorins, play pivotal roles in modulating spinal cord circuit formation, maintenance, and function (59, 63, 64). Indeed, neuronal subtypes that produce ligands, such as neuropeptides, may modulate the function of subtypes expressing their receptors (65). Yet relatively little is known about ligand-receptor (L-R) interaction profiles in the spinal cord at the level of neuronal subtype. To this end, we queried expression patterns across cell subtypes for known neurotransmitters, neuropeptides, neurotrophins, and semaphorins and their receptors, as well as other L-R pairs, and used NICHES (66) to infer intercellular signaling based on the expression levels of the L-R pairs. Our analysis uncovered ligand expression patterns of putative sender cells and receptor expression patterns of putative receiver cells within the dorsal horn (Figures 5A and S5).

**Figure 5.**
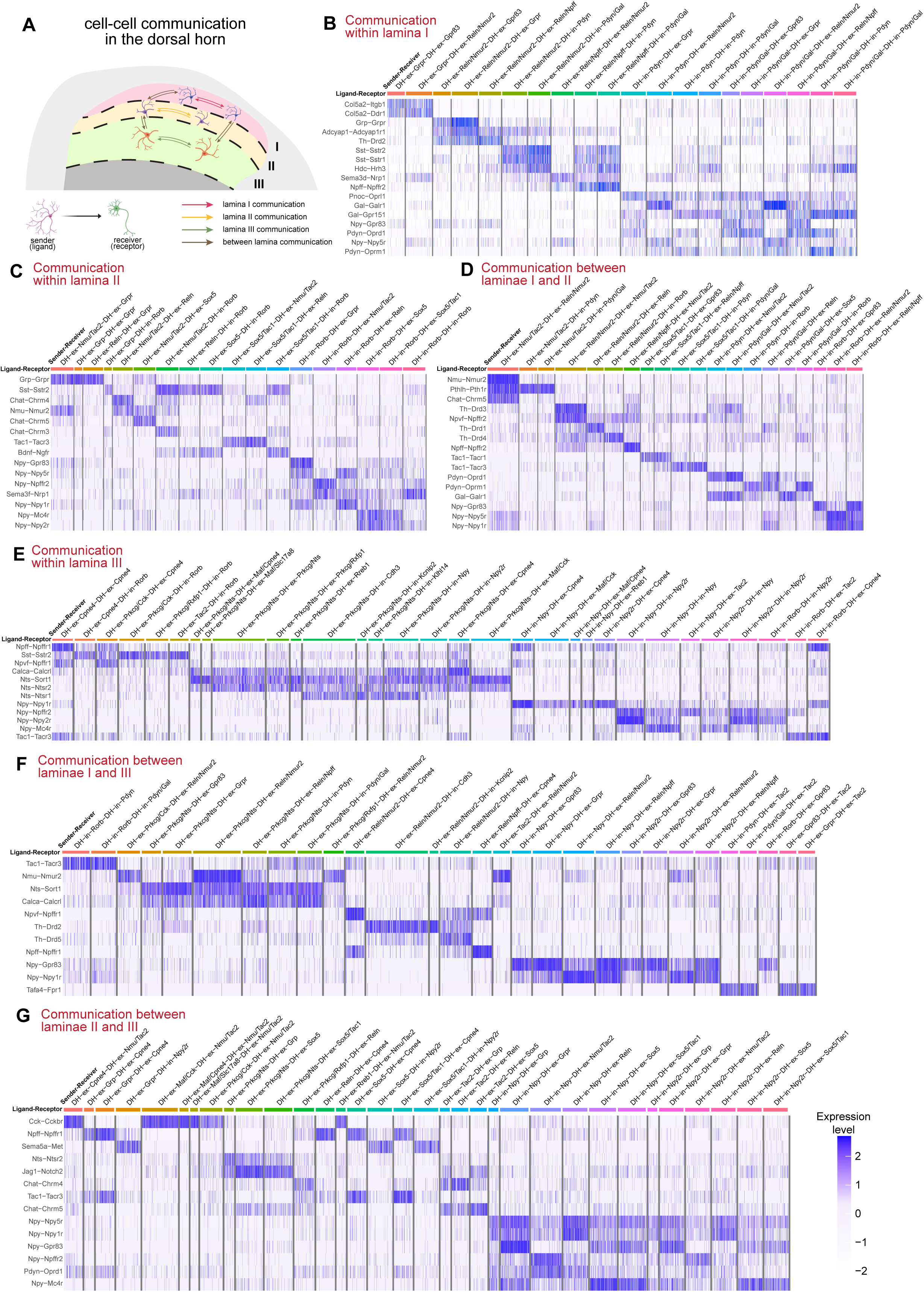
Cell-cell communication through ligand-receptor interactions between dorsal horn neuronal subtypes. (A) Schematic overview of cellular communication analyzed in the dorsal horn. Illustration summarizing the spatial domains of laminae I–III in the dorsal horn and the directionality of cell–cell communication events inferred from ligand–receptor expression profiles. Communications within each lamina and across laminae are represented with arrows. (B, C, and E) Communication of neuron populations within lamina I (B), II (C), and III (E). (D, F, and G) Communication of neuron populations between laminae I and II (D), I and III (F), II and III (G). The color scale indicates normalized expression levels of the ligand–receptor pair.

We observed many neuropeptides and their cognate receptors that were differentially present between putative sender and receiver subtypes within individual dorsal horn lamina (Figures 5B, 5C, 5E, S5A, and S5B). For example, within lamina I (Figure 5B), DH-in-Pdyn/Gal neurons, marked by the expression of *Gal* encoding the peptide Galanin (67), may communicate with DH-ex-Reln/Nmur2 neurons, which express galanin receptor type 1 (*Galr1*). DH-ex-Reln/Npff neurons may signal to DH-in-Pdyn/Gal neurons via neuropeptide FF (*Npff*)–neuropeptide FF receptor 2 (*Npffr2*), a neuropeptide pathway that modulates hyperalgesia and opioid sensitivity (Figure 5B) (68). Such reciprocal interaction patterns between neuron subtypes of the *Reln* and *Pdyn* families through different neuropeptides featured both pro-and anti-nociception signaling. Within lamina II (Figure 5C), expression of *Grp*, which encodes gastrin-releasing peptide associated with itch transmission (69), was detected among DH-ex-Nmu/Tac2, DH-ex-Grp, and DH-ex-Reln sender subtypes. DH-ex-Grpr vertical neurons were identified as putative receivers, consistent with their elaborate ventral orientation of dendrites into the dorsal horn (41). Moreover, neuropeptide Y (*Npy*)–neuropeptide Y receptor type 2 (*Npy2r*) communication was primarily observed between DH-in-Rorb sender neurons and DH-ex-Sox5 and DH-ex-Sox5/Tac1 receiver neurons (Figure 5C). Among neuronal subtypes residing in lamina III, neuropeptide Y–neuropeptide Y receptor type 1 (*Npy1r*) interaction, which modulates light touch and mechanical itch transmission (70), was observed between DH-in-Npy sender neurons and many receiver subtypes, including DH-ex-Cpne4, DH-ex-Maf/Cck, DH-ex-Maf/Cpne4 and DH-ex-Rreb1 neurons (Figure 5E). In addition, several neuronal subtypes expressing *Npff* might communicate with DH-ex-Cpne4 neurons through *Npffr1* (neuropeptide FF receptor 1), a pro-nociceptive L-R interaction that shapes pain sensitivity (Figure 5E). Similarly, DH-ex-Prkcg/Nts neurons express *Nts* (neurotensin), which may act on a number of inhibitory neuron subtypes expressing *Ntsr1* (neurotensin receptor type 1) (71), consistent with the widespread synaptic connections of Prkcg^+^ neurons in lamina III (Figure 5E) (7).

Some L-R interactions were consistently detected between neuronal subtypes within the same lamina (Figures 5B, 5C, and 5E). For instance, somatostatin signaling, which inhibits nociception (72), was detected between the *Reln* family neurons (i.e., DH-ex-Reln/Nmur2 and DH-ex-Reln/Npff) as senders and the *Pdyn* family neurons as receivers (Figure 5B). Within laminae II and III, potential somatostatin signaling was also observed between multiple senders, including DH-ex-Nmu/Tac2, DH-ex-Reln, and DH-ex-Prkcg/Cck neurons, and the receiver DH-in-Rorb and DH-ex-Cpne4 neurons (Figures 5C and 5E).

Our analysis also revealed a plethora of potential L-R interactions mediating CCC between sender and receiver neurons located in different laminae (Figures 5D, 5F, 5G, and S5C-S5E). As expected, Neuromedin U signaling mediated by the *Nmu*-*Nmur2* pair (73), was detected between DH-ex-Nmu/Tac2 senders in lamina II and DH-ex-Reln/Nmur2 receivers in lamina I (Figure 5D). We observed that one neuronal subtype may serve as a sender subtype to multiple subtypes. For example, strong neurotensin L-R interaction (Nts-Sort1) was detected between DH-ex-Prkcg/Nts as a sender cell type and multiple subtypes in the superficial lamina as receiver cell types (Figure 5F), consistent with the presence of synaptic connections in Prkcg^+^ neurons in the superficial laminae (7). In some cases, multiple subtypes in one lamina could serve as senders to a single receiver subtype in another lamina. For example, DH-ex-Nmu/Tac2 receiver neurons in lamina II expressed *Cckbr* (cholecystokinin B receptor), and multiple neuronal subtypes in lamina III were detected as cholecystokinin senders (Figure 5G), including DH-ex-Cpne4 and all three excitatory neuronal subtypes of the *Maf* family. This convergence of cholecystokinin signaling from multiple senders onto DH-ex-Nmu/Tac2 receiver neurons suggests that DH-ex-Nmu/Tac2 neurons may be an important neuronal subtype mediating cholecystokinin’s contribution to pain. We further performed the CCC analyses using dorsal-intermediate, dorsal-ventral, and intermediate-ventral neuronal subtypes, which reveal a broader view of candidate neuromodulatory CCC that may be involved in sensory, autonomic, and motor circuits (Figures S6A-S6C).

We also investigated potential L-R interactions between neuronal and non-neuronal cells within individual lamina (Figures S6D-S6F). In contrast to the prevalence of neuropeptide signaling detected between neuronal subtypes, many of the top L-R interactions for neuronal–non-neuronal interactions belong to semaphorin and notch signaling pathways. Intriguingly, *Adcyap1*, which encodes adenylate cyclase-activating polypeptide 1 (PACAP), was detected across multiple sender neuronal subtypes of lamina III, whereas *Adcyap1r1*, encoding the PACAP receptor, was most enriched in astrocytes as receivers (Figure S6F), suggesting region-specific L-R interactions between neurons and astrocytes. Collectively, these findings reveal a large variety of candidate CCC patterns within and between dorsal horn laminae of the spinal cord. Each lamina exhibits both unique and shared L-R interactions among excitatory and inhibitory microcircuits, defining distinct lamina-specific CCC motifs.

### Identification of spinal neuron subtypes and neuropeptide signals associated with neuropathic pain

The spinal cord is a critical hub for processing nociceptive signals, integrating sensory input from the body and modulating these signals through complex neural circuits that govern local reflexes and output to the brain (3, 4). Within the framework of the gate control theory, a balance of excitation and inhibition among spinal cord neurons may act as the neural basis of a “gate”, a dynamic checkpoint by which the relative strengths of excitatory and inhibitory inputs control the transmission of nociceptive signals to the brain (65). Excitatory interneurons amplify nociceptive inputs, while inhibitory interneurons suppress them using the neurotransmitters GABA and glycine (65, 74). Disruption of spinal cord circuit elements can lead to abnormal processing of nociceptive and innocuous sensory inputs, contributing to heightened pain sensitivity and chronic pain. Transcriptional changes in the spinal cord have been associated with different pain states (26, 75); however, the spatial and cell-type-specific resolution of these changes remains poorly understood. The MERFISH platform enables the identification of spinal neuron subtypes and the transcriptomic alterations that may underlie spinal cord circuit dysfunction and lead to neuropathic pain.

To this end, we employed the spared nerve injury (SNI) model of neuropathic pain (76), in which two of the three branches of the sciatic nerve are transected, leaving the sural branch intact (Figure 6A). Sham mice, having their sciatic nerve exposed but not transected, served as controls. The dynamic brush assay was used to confirm the development of robust mechanical allodynia in SNI animals that persisted for at least 7 days post-surgery (Figure 6B). Lumbar spinal cords were collected from both sham and SNI mice for MERFISH profiling to assess transcriptional alterations associated with neuropathic pain (Figure 6A). Overlap of UMAP visualizations affirmed the convergence of sham and SNI datasets, ensuring a reliable comparison for revealing the effects of neuropathic pain (Figure S7A).

**Figure 6.**
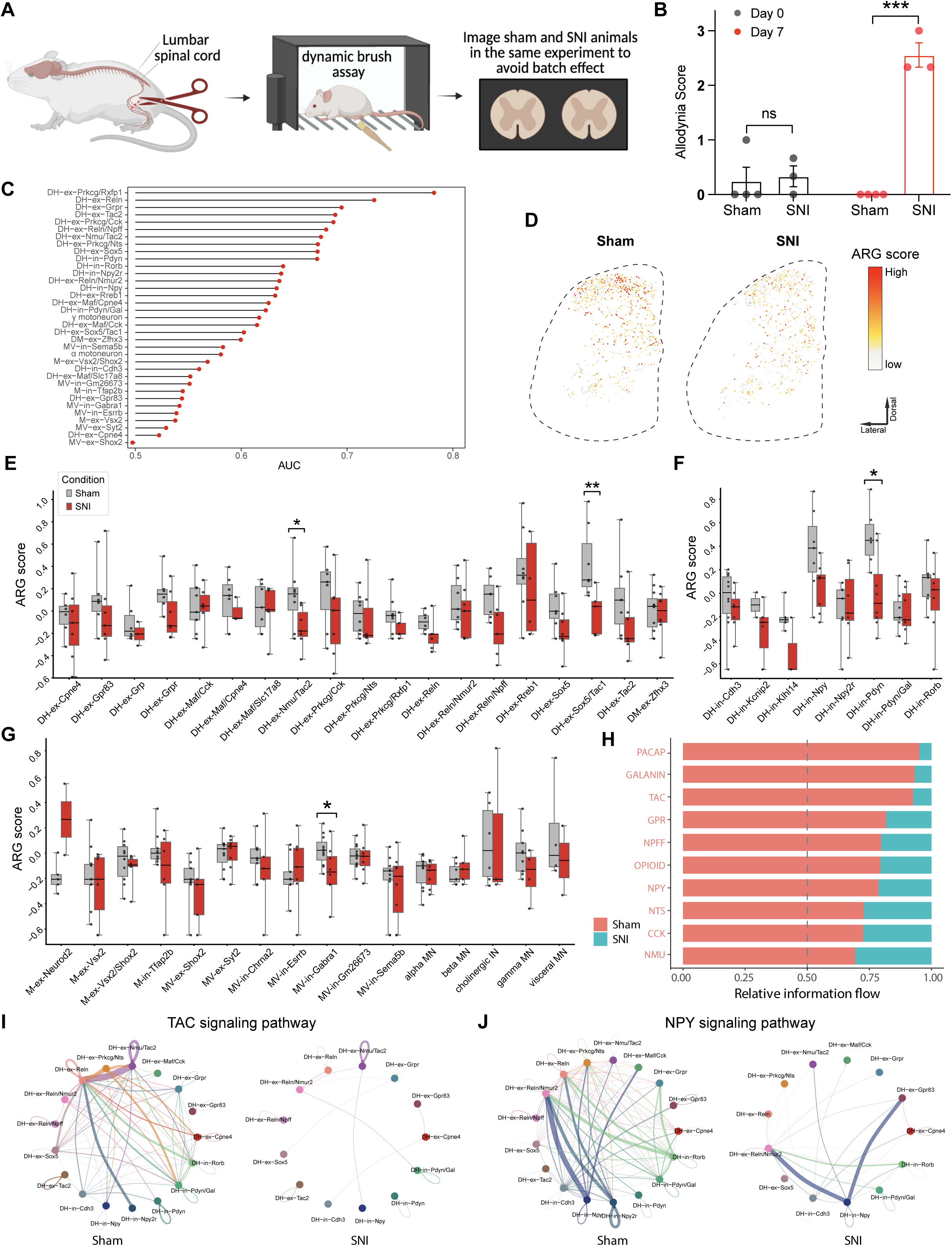
Nerve injury that causes neuropathic pain leads to molecular and cellular changes in spinal cord neurons. (A) Overview of the spared nerve injury (SNI) sample preparation as a model of neuropathic pain. For each MERFISH experiment, spinal cord slices from each axial level of each of the sham and SNI conditions are imaged together to avoid batch effects. (B) Quantifications of allodynia scores at day 0 (baseline) and day 7 post-surgery showed a significant increase in brush-evoked responses in the SNI group. ns, not significant; ∗∗∗p < 0.001; error bars indicate SEM. (C) Transcriptionally perturbed neuron populations predicted by Augur and ranked by AUC value. AUC shows the area under the ROC curve of the predictions. (D) Spatial distribution of neurons colored by activity-related gene (ARG) scores in sham (gray) and SNI (red) animals on representative sections. The dashed line delineates the contour of the left lumbar spinal cord. (E-G) Box plots of ARG scores across neuronal subtypes in sham and SNI animals. Wilcoxon rank-sum tests. *p < 0.05, **p < 0.01. (H) Significant neuropeptide signaling pathways were ranked according to differences in overall information flow, defined as the sum of communication probabilities in a given inferred network. Those colored red are more enriched in the sham spinal cords. (I and J) Detected possible interactions in the tachykinin (I) and NPY (J) signaling pathways between pairs of dorsal horn neuronal subtypes. Edge width represents the communication probability, and each edge is colored to match its sender neuronal subtype, indicating the direction of signal origin.

To identify cell types in the ipsilateral lumbar spinal cord that are most transcriptionally altered during the neuropathic pain state, we employed Augur (77), an algorithm that can rank the cell types most responsive to biological perturbations based on transcriptomic changes. Augur determines an area under the curve (AUC) score to quantify how accurately individual cell types can be distinguished between conditions (77). In general, neuronal populations in the dorsal horn were more transcriptionally perturbed with higher AUC scores after SNI compared to other spinal cord regions. Indeed, the top nine most perturbed subtypes were all dorsal horn excitatory neurons (Figure 6C). Among these, DH-ex-Prkcg/Rxfp1 and DH-ex-Reln neurons were most strongly perturbed, followed by DH-ex-Grpr, DH-ex-Tac2, DH-ex-Prkcg/Cck, DH-ex-Reln/Npff, DH-ex-Nmu/Tac2, DH-ex-Prkcg/Nts, and DH-ex-Sox5 neurons. The most affected inhibitory subtypes were DH-in-Pdyn, DH-in-Rorb, and DH-in-Npy2r neurons. To address how the expression of specific genes is impacted in these subtypes, we calculated feature importance genes that contributed most to the transcriptional perturbation in these top subtypes (Figures S7B and S7C). This analysis revealed several new genes that are impacted within each neuronal subtype in SNI, such as the reduction in the *Qdpr* and *Sema5b* genes in the DH-in-Pdyn neurons (Figure S7C). These findings indicate that neuropathic pain is associated with transcriptional alterations in select dorsal excitatory neuron subtypes and, to a lesser extent, subtypes of dorsal horn inhibitory neurons.

Neuropathic pain is often associated with altered spinal cord activity, including decorrelated dorsal horn network activity (27). Therefore, we next sought to identify cell types whose activity may be altered in the neuropathic pain state by asking whether SNI is associated with changes in the expression levels of activity-regulated genes (ARGs), including *Arc*, *Junb*, *Fos*, *Fosb*, *Npas4*, and *Nr4a1*. We calculated the ARG score for each neuronal subtype and compared them between the sham and SNI groups. Spatial visualization of ARG scores across representative lumbar sections revealed a reduction in ARG expression as a proxy for reduced neuronal activity in SNI mice compared to sham mice (Figure 6D). Of the four neuronal subtypes that exhibited significant decreases in ARG scores in the SNI group, DH-ex-Nmu/Tac2, DH-ex-Sox5/Tac1, and DH-in-Pdyn neurons are located in the dorsal horn, whereas MV-in-Gabra1 neurons reside in the intermediate zone and ventral horn (Figures 6E-6G). Meanwhile, ARG for parvalbumin (PV)-expressing inhibitory interneurons did not change significantly compared to sham animals (Figure S8I), suggesting that PV+ interneurons do not exhibit detectable transcriptional changes in activity-regulated genes in this model of neuropathic pain (78). This reduction in ARG scores in several subtypes suggests that the neuropathic pain state elicits hypoactivity within defined dorsal horn subtypes that are associated with processing nociceptive and innocuous signals.

To assess how cell-cell communication between spinal cord neuron subtypes is impacted in neuropathic pain, we employed CellChat (79) to investigate inferred neuropeptide signaling interactions among dorsal horn neuronal subtypes in the ipsilateral lumbar spinal cords. We quantified the information flow, which is defined by the sum of communication probabilities among all pairs of neuronal subtypes in the inferred network in sham and SNI spinal cords (Figure 6H). We observed many expected neuropeptide signaling between neuronal subtypes in the sham spinal cord, including the tachykinin and NPY signals released from DH-ex-Nmu/Tac2 and DH-in-Npy subtypes, respectively (Figures 6I and 6J). Several peptide L-R motifs, including those for PACAP, galanin, opioid, and NPY, exhibited decreased overall information flow in the SNI condition compared to the sham condition (Figures 6H, S8A-S8H). For example, tachykinin and NPY L-R communication probabilities were decreased both between subtypes and within the same subtypes in SNI animals (Figures 6I and 6J). Galanin was detected between DH-in-Pdyn neurons as senders and five other subtypes in the dorsal horn (67), and its inferred signaling from the DH-in-Pdyn neurons was only detected in DH-ex-Reln/Nmur2 in the SNI condition (Figure S8B). Consistent with previous observations that peripheral axonal injury reduces opioid action in the spinal cord (80), our results further reveal that the reduction was more prominent in certain opioid sender subtypes, such as Pdyn subtypes (Figure S8G). PACAP, which is broadly detected in the Reln family neurons as senders in the sham spinal cord, is detected at a low level in the SNI condition (Figure S8H). Together, these findings reveal a broad attenuation of autocrine and paracrine neuropeptide signaling between selective sender-receiver neuronal subtypes during the neuropathic pain state.

## Discussion

In this study, we used MERFISH to gain insight into the molecular and cellular organization of the mouse spinal cord. By profiling cells across multiple axial levels in both sexes, we resolved the cellular landscape of neurons along both the R-C and D-V axes and revealed sex differences in the spinal cord. We also identified dorsal horn neuronal subtypes that undergo transcriptional and neuropeptide signaling alterations associated with neuropathic pain. These data provide a resource and computational framework for systematic mapping of molecules, cells, and their organizational properties. Given the recent growing literature on single-cell and spatial transcriptomic atlases of the spinal cord (18, 51, 81), systematic cross-dataset integration will be important for building toward a harmonized taxonomy of spinal cord cell types. To enhance accessibility of these datasets, we developed a web-based browser for the spatial visualization of our data (https://github.com/shanmeltzer/mSpinalCord_MERFISH).

The spinal cord contains an intricate network of interconnected neurons that integrate somatosensory inputs and coordinate autonomic and motor outputs (3). Many neuronal subtypes are present in different relative abundances across axial levels, suggesting they may be involved in body region-specific sensory and motor circuit functions. For example, DH-ex-Gpr83 neurons (34), a subtype likely to be superficial-lamina projection neurons, are most abundant in the cervical level, followed by the lumbar level, suggesting that they may be more engaged in relaying limb-related sensory information to the brain (82). Similarly, neuronal subtypes with the highest percentages at sacral levels (Figure 2B) may be more engaged in pelvic organ control, sexual function, and tail reflexes (20). These findings support the idea that spinal cord function is regionally diversified through variation in the abundance of neuronal subtypes across different axial levels.

Although sex differences in the number of certain neuronal subtypes were previously observed in the spinal cord (55, 58), a comprehensive analysis of sex differences of spinal cell types across axial levels had not been performed. Even though previous analyses reported sex differences in the number of spinal cord visceral motoneurons between males and females (55, 58), our findings show that this difference is primarily driven by higher numbers of these neurons in the lumbar level of males. Furthermore, sexual dimorphism in the percentage of dorsal horn subtypes observed here may contribute to the sex differences in touch and pain sensitivity, while differences in the intermediate and ventral spinal cord may reflect differential engagement of these subtypes in sensorimotor and autonomic spinal circuitry (83). Future studies dissecting these sex differences at the molecular and circuit levels will help elucidate how they may influence spinal cord physiology and the neural control of bodily functions.

Our results provide a resource for future investigations into how local circuit motifs and intercellular signaling modules underlie or shape spinal cord circuit function. Our cell-cell proximity analysis revealed that the somas of many cell-type pairs are closely positioned to one another in space. Future connectomic and electrophysiological analyses are needed to test whether functional connectivity indeed exists between the spatially proximate subtypes described here. Our CCC analysis based on L-R gene expression also identified a range of putative L-R interactions between neuronal subtypes within and between laminae of the dorsal spinal cord. These findings support the view that the spinal cord operates as a mosaic of microcircuits, wherein the cellular neighborhood composition and L-R interactions enable functional coupling above and beyond fast excitatory and inhibitory transmission. Future validations and functional analyses of these L-R interactions at the level of cell subtypes will reveal whether and how these inferred CCC interactions may shape somatosensory information processing.

Identifying spinal circuit motifs and loci of dysfunction underlying central sensitization and allodynia following various types of nerve injury represents an active area of research (4, 26, 38). We observed most transcriptomic perturbations across select excitatory neuron subtypes within the dorsal horn, and decreased ARG scores in three neuronal populations in the dorsal horn and one inhibitory subtype in the intermediate zone. The ARG score combines six activity-regulated genes that are rapidly induced and peak within 30 min of neuronal activity (53), with varying timeframes of returning to baseline (53). By combining genes with distinct decay kinetics into a composite score, our approach integrates neuronal activity over a broader time window (84). These transcriptomic changes may be caused by alterations in behavioral state, sensory activity, or postural changes that may occur in the neuropathic pain state. A recent study reported increased activity-regulated gene expression in SNI mice across other populations not observed in our analysis, and these different outcomes may reflect differences in the timing post-injury and the neuronal activation state during tissue harvesting (4, 26).

Our observation of a decreased ARG score in DH-in-Pdyn neurons is consistent with previous work demonstrating that Pdyn/Dyn lineage inhibitory neurons engage in circuit-gating protective reflexes by A-LTMRs and gate mechanical pain (39, 40). Our observation that DH-in-Pdyn neurons are the only inhibitory subtype with a reduced ARG score supports the idea that targeting the activity of this inhibitory neuron subtype may provide a strategy for treating neuropathic pain. Moreover, the decreased ARG scores across several excitatory neuron types point to a possible reduction in the excitatory drive of inhibitory interneurons and possible disinhibition of spinothalamic tract projection neurons (59). We did not observe significant changes in the ARG scores in the Parvalbumin neurons that gate mechanical inputs in neuropathic pain (Figure S8I), suggesting their gate-keeping role in pain is independent of the ARG expression levels (78, 85). Furthermore, the subtype-resolved alterations in neuropeptide signaling identified specific ligand-receptor interactions as testable targets for future functional and therapeutic studies aimed at alleviating neuropathic pain. Our SNI collection time point primarily captures the acute phase of neuropathic pain, and longitudinal transcriptomic profiling encompassing both the initiation and chronic phases will reveal the temporal dynamics of the transcriptomic transitions during the “chronification” of pain. Furthermore, whether any sexually dimorphic transcriptomic responses emerge following SNI remains to be determined. Last, it is possible that in pathological pain conditions, spinal neuronal subtypes show reduced activity in an unstimulated resting state, while showing increased activity after sensory stimulation, such as the Grpr^+^ neurons in diabetic polyneuropathic pain (86). Hence, future studies utilizing a variety of somatosensory stimuli will provide insights into the neuronal subtypes differentially impacted by sensory stimuli in neuropathic pain. Together, these findings provide a basis for future functional analyses of circuit-level dysfunction associated with neuropathic pain states.

## Materials and Methods

Adult female and male C57BL6 wildtype mice, spared nerve injury (SNI) mice, and sham-surgery mice aged 8-10 weeks were used in the study. Animals were perfused, followed by spinal cord collection and sectioning for MERFISH imaging. MERFISH data were analyzed by customized analysis pipelines. Detailed descriptions of mouse strains, MERFISH panel design and imaging, RNAscope, immunohistochemistry, and data analysis pipelines are provided in the SI Appendix.

## Data and code availability

MERFISH data have been deposited in Zenodo (https://zenodo.org/records/20533289). All original analysis code is available at https://github.com/shanmeltzer/mSpinalCord_MERFISH.

## Acknowledgements

We thank members of the Meltzer and Ginty lab and Dr. Christos Constantinidis for their feedback on the manuscript. We thank Dr. Benjamin Brown for helping with the relative depth analysis. This work was supported by the HHMI Hanna Gray Fellowship (S.M.), Edward Mallinckrodt, Jr. Foundation (S.M.), Sloan Research Fellowship (S.M.), NIGMS MIRA R35 GM146960 (X.M.Z), NSF CAREER DBI-2541736 (X.M.Z.), Vanderbilt Seeding Success Grant FF_300627 (S.M. and X.M.Z), Vanderbilt CCSB Accelerator Fund (S.M. and X.M.Z), NIH-NIGMS-T32-GM149363-02 (M.S.), NIH 1R01 AT011447-01 (D.D.G.), NIH R35 5R35NS097344-05 (D.D.G.), and NIH R35 5R35NS132196-02 (D.D.G.). D.D.G. is an investigator of the Howard Hughes Medical Institute. This article is subject to HHMI’s open access to publications policy. HHMI lab heads have previously granted a nonexclusive CC BY 4.0 license to the public and a sublicensable license to HHMI in their research articles. Pursuant to those licenses, the author-accepted manuscript of this article can be made freely available under a CC BY 4.0 license immediately upon publication.

## Author contributions

S.M. and D.D.G. conceived the study. S.M., Q.W., E.O.-A., M.S., and G.C. performed the MERFISH and RNAscope experiments. J.P. and B.S. generated the SNI mouse model and performed the behavior tests. S.M., Q.W., Y.H., Y.Z., X.M.Z. analyzed the data or supervised the experimental data analysis. Q.W. and Y.H. developed the data processing and analysis pipeline. Q.W. constructed the data web browsers. S.M., Q.W., Y.H., and X.M.Z. wrote the manuscript with inputs from all authors.

