## supplemental information for "Spatial transcriptomics reveals organizational properties of the mouse spinal cord and alterations in neuropathic pain"

PNAS

**This PDF file includes:**

- Supplementary Information for Materials and Methods
- Figures S1 to S8
- Tables S1 and S2
- Supplemental Information References

**Supplementary Information for Materials and Methods**

**Animals**

Animal care and experiments were conducted in accordance with the National Institutes of Health (NIH) guidelines and were approved by the Institutional Animal Care and Use Committee (IACUC) of Harvard Medical School and Vanderbilt University. Adult female and male C57BL/6 mice of 8-10 weeks old were used in the study. Mice were maintained on a 12-hour light/12-hour dark cycle with ad libitum access to food and water.

**Immunohistochemistry and confocal imaging**

Spinal cord slides were allowed to dry on the benchtop overnight and stored at -20 °C for long-term storage. Slides were defrosted at room temperature for 20 minutes, and then a hydrophobic barrier was applied using a PAP pen (H-4000, VWR/Vector Lab) around the sections. Then, sections were rehydrated with filtered 1xPBS for 3 × 5 minutes, blocked for one hour at room temperature with 1xPBS with 0.1% Triton X-100 and 5% normal goat or donkey serum (Jackson Immuno, 005-000-121). Sections were then incubated with primary antibodies in blocking solution (5% serum in PBS with 0.1% Triton X-100) overnight at 4 °C. Sections were washed 4x5 minutes using 1xPBS with 0.02% Tween-20 and incubated with secondary antibodies in blocking solution (5% serum in PBS with 0.1% Triton X-100) at room temperature for 2 hours. After secondary antibody incubation, sections were washed 4x5 minutes with 1xPBS with 0.02% Tween-20 detergent and mounted with fluoromount-G (Southern Biotech). All slides were stored at 4 °C for up to one month until imaged. Z-stack images of the entire spinal cord sections were obtained on the Zeiss LSM900 confocal microscope with a 20x objective, and the tile-stitching function was used. Images were analyzed and prepared in ImageJ software.

The following primary antibodies were used: Isolectin B4 647 (Invitrogen, RRID: SCR_014365, 1:500), rabbit anti-CGRP (Immunostar, Cat#24112, RRID: AB_572217, 1:500), and mouse anti-Kv4.3 (Antibodies Incorporated, Cat# 75-017, RRID: AB_2131966, 1:500). Secondary antibodies include Alexa 488 and 546 conjugated goat anti-mouse and goat anti-rabbit antibodies at a 1:500 dilution (Life Technologies).

**RNAScope**

Adult female and male C57BL6 mice of 8-10 weeks old were euthanized by $CO_2$ followed by decapitation. Spinal cords were dissected out, and lumbar segments were quickly frozen in dry-ice-cooled 2-methylbutane, stored at -80 °C, and then sectioned at a thickness of 16-20 μm for RNA transcript detection by RNAscope (Advanced Cell Diagnostics) following the manufacturer's protocol. For the γ motoneurons, we modified the sample pre-treatment by increasing the fixation time to 1.5 hours and decreasing permeabilization time to 15 minutes. These probes were used: Mm-Chat (Cat# 408731), Mm-Nos1-C2 (Cat# 437651-C2), Mm-Esrrg-C2 (Cat# 495121-C2), and Mm-Npas1-C3 (Cat# 468851-C3).

**MERFISH gene panel selection and probe construction**

A panel of 500 genes (Supplementary Table 1) that contained 4 categories of genes: (1) cell-type markers; (2) neuronal function regulatory genes; (3) ligand-receptor pair genes, and (4) neuronal activity-regulated genes (ARGs) was designed. For cell-type marker genes, 94 neuronal markers for subtypes of excitatory, inhibitory, and cholinergic neurons in the spinal cord, and Neurod1 and Rbfox3 as general neuronal markers were selected. 29 non-neuronal markers were also included for identifying other major cell types (i.e., astrocytes, microglia, oligodendrocyte precursor cells, oligodendrocyte progenitors, oligodendrocytes, endothelial cells, ependymal cells, pericytes, meningeal cells, Schwann cells, and peripheral glia). These genes were obtained as top-enriched cluster markers in a published harmonized atlas of mouse spinal cord cell types (1, 2). Neuronal function regulatory genes comprised genes encoding neurotransmitters and their receptor channels, and neuroactive peptides and their receptors. Genes that encoded ligand-receptors (semaphorins, neurotrophins, and Notch) and cell adhesion molecules (clustered protocadherins) were included as well. In addition, we added to the panel neuronal activity-regulated genes (ARGs) to report the activity history of spinal cord neurons (2, 3), resulting in a total of 500 genes. All genes in the panel were included for subsequent analyses. MERFISH encoding probes were designed and synthesized by Vizgen using a commercial pipeline.

**MERFISH tissue collection, processing, and imaging**

Mice were anesthetized with isoflurane and perfused with 4% PFA. Spinal columns were dissected and post-fixed in 4% PFA for 1.5 hours at 4°C and rinsed with 1X PBS (made with nuclease-free water). Spinal cords were then dissected out and cryoprotected in 30% sucrose in 1X PBS with RNase inhibitor (NEB, Cat# M0314L) overnight at 4°C. The spinal cords were embedded in OCT on a mixture of dry ice and ethanol bath, and stored at −80 °C until sectioning. The spinal cord tissues were sectioned at different thicknesses (10 μm and 60 μm). 60-μm-thick sections were collected for RNA extraction with the Quick-RNA FFPE Miniprep kit (Zymo, Cat# R1008) to assess RNA quality of each animal. Average DV200 values were 84, with the lowest DV200 value being 78.82. Sections at 4 different spinal axial levels (cervical, thoracic, lumbar, and sacral) were picked for MERFISH from serial 10-μm-thick sections, with the adjacent sections prepared for immunohistochemistry and confocal imaging.

Standard MERFISH protocol was performed according to the manufacturer's protocol (Vizgen, fixed frozen sample preparation user guide). Briefly, spinal cord sections were mounted onto a MERSCOPE slide (Vizgen, REF. PN 20400001) and kept at −20 °C for 5 minutes to allow for tissue adherence to the slide. Sections were washed with 1X PBS and stored in 70% ethanol at 4 °C overnight to allow permeabilization of the tissue. After permeabilization, sections were washed with Sample Prep Wash Buffer (Vizgen, REF. PN 20300001), followed by Formamide Wash Buffer (Vizgen, REF. PN 20300002), drained, and then hybridized with 50 μL MERSCOPE gene panel mix containing customized encoding probes. The slide was covered with Parafilm in a sealed Petri dish and incubated at 37 °C for 36 to 48 hours. Later, sections were washed twice with Formamide Wash Buffer, followed by Sample Prep Wash Buffer, and drained. A Gel Coverslip (Vizgen, REF. PN 30200004) coated with Gel Slick Solution (Lonza, Cat# 50640) was placed onto the sections that were covered with 50 μL mixture of Gel Embedding Premix (Vizgen, REF. PN 20300004), 10% ammonium persulfate solution and N,N,N',N'-tetramethylethylenediamine, and incubated at room temperature for 1.5 hours to allow embedding of the sections into a thin layer of poly-acrylamide gel. After removal of the coverslip, sections were incubated at 37°C with 5 mL solution containing warmed Clearing Premix (Vizgen, REF. PN 20300003) and Proteinase K (NEB, Cat# P8107S) for at least 24 hours to allow clearing of the tissue, preparing for MERFISH imaging.

MERFISH imaging was performed on a MERSCOPE instrument following the manufacturer's protocol (Vizgen, MERSCOPE instrument user guide). Briefly, a MERSCOPE Imaging Cartridge (Vizgen, REF. PN 20300019) was activated by adding a mixture of 250 µL Imaging Buffer Activator (Vizgen, REF. PN 20300022) and 100 µL RNase inhibitor and covered with 15 mL mineral oil. The cartridge was loaded into the MERSCOPE instrument, and the fluidics system was primed according to instructions. The section slide was taken from the clearing solution and washed twice with Sample Prep Wash Buffer before staining with DAPI and PolyT Staining Reagent (Vizgen, REF. PN 20300021) for 15 minutes. After staining, the slide was assembled to the MERFISH Flow Chamber, and the chamber was loaded into the instrument and connected to the fluidics system. Low-resolution mosaics were acquired initially to guide the selection of regions of interest, which were subsequently imaged with a 60x objective at 7 planes on the z-axis.

**MERFISH image processing and data preprocessing**

Images were decoded by MERSCOPE Instrument Software v232, which generated detected transcripts, cell-by-gene, and cell-metadata matrices. Cell segmentation, transcript partitioning, and cell metadata calculation were performed using the Vizgen Post-processing Tool (VPT, v1.3.0) with the Cellpose2 plugin (4, 5) (vpt-plugin-cellpose2, v1.0.1), where a standard "nuclei" model was applied using the DAPI channel to delineate cell boundaries. After segmentation, transcripts were partitioned into cells, and cell metadata was calculated.

The data preprocessing pipeline was implemented in Python with anndata (v0.7.5.3) (6) and Scanpy (v1.9.1) (7) packages. For each MERFISH experiment, an anndata object was created by combining gene expression data and cell metadata. To account for samples and experiments associated with captured cells, additional metadata, including mouse ID, sex, section ID, spinal axial level, and experiment batch were added. To remove low-quality captures, cells with total transcripts less than 25 were removed, and genes labeled as "blank" were removed as well. AnnData objects were subsequently loaded to the MERSCOPE Visualizer for manual inspection. After excluding cells of crescent shapes reflecting fragments of fiducial beads, and cells falling outside of any defined tissue section, filtered objects were concatenated into a single dataset. To standardize gene expression values, total RNA counts per cell were normalized (scanpy.pp.normalize_total), log-transformed (scanpy.pp.log1p), and converted to Z-scores (scanpy.pp.scale). To denoise the data, principal component analysis (PCA) was performed for dimensionality reduction (scanpy.pp.pca). A k-nearest neighbors (KNN) graph was constructed using scanpy.pp.neighbors (n_neighbors=10, n_pcs=20), which was then used to compute a Uniform Manifold Approximation and Projection (UMAP) (8) embedding for visualization (scanpy.tl.umap).

**Cell clustering, cluster refinement, and manual annotation**

We leveraged a multi-tier clustering strategy with the Leiden community detection algorithm (9) to achieve high-resolution identification of cell populations in our dataset. Initial clustering was performed using scanpy.tl.leiden (resolution=0.5) function in Scanpy*.* 14 clusters of conventional cell classes were determined and annotated (i.e., five clusters of neurons, three clusters of oligodendrocyte lineage, two clusters of astrocytes, one cluster of endothelial cells, meningeal cells, microglia, and peripheral cells) based on the gene expression of cell class markers derived from the reference atlas (1), as well as the spatial locations of cells.

To further reveal fine-grained cell types and subtypes, cells from each cluster were individually subsetted from the whole dataset and reanalyzed following the same procedure as described in the initial clustering. Briefly, for each subset, PCA, KNN graph, and UMAP were recomputed and reconstructed, followed by Leiden clustering. Three clusters were excluded from subclustering after manual review, as their expression patterns of marker genes were indicative of homogeneous populations corresponding to astrocytes-1, oligodendrocyte-2, and microglia of the atlas cell types, respectively. We named the neuronal subtypes using a 3-element system (location-ex/in-marker). In the case where a recognized family marker gene is expressed across multiple subtypes, the family marker was affixed before the individual marker (e.g., Prkcg/Nts for Prkcg as family marker and Nts as individual subtype marker). The resulting subclusters were annotated based on expression patterns of marker genes to best match the cell types and subtypes reported in the atlas (1).

**Assessment of cell type concordance between manual and transferred labels**

We applied a reference-based label transfer approach to annotate cell types in the MERFISH dataset based on a snRNA-seq reference (10) using SingleR (v3.20) (11). For both datasets, the raw count matrices were extracted and transposed to conform to the standard gene-by-cell orientation. Seurat objects were constructed using Seurat (v4.1.1) (12) with the transposed matrices and corresponding metadata. Counts were normalized with LogNormalize to generate comparable expression values across cells, and normalized Seurat objects were converted to SingleCellExperiment (v1.16.0) objects to ensure compatibility with SingleR. The predefined cell-type annotations in the snRNA-seq dataset were used as reference labels in SingleR, which was run with the snRNA-seq dataset as reference and the MERFISH dataset as query. Cell type assignment was based on marker genes identified by Wilcoxon rank-sum tests (de.method = "wilcox"), selecting the top 10 differentially expressed genes per cluster (de.n = 10). The output provided predicted cell-type labels for each MERFISH cell based on transcriptional similarity to the snRNA-seq reference.

To assess the accuracy of reference-based label transfer, we compared SingleR-predicted labels with our manual annotations of the queried MERFISH dataset by computing a confusion matrix between predicted and manual labels and quantifying the proportion of overlapped cells for each category. For each MERFISH cell type, the percentage of cells with the predicted label matching the manual annotation was calculated to generate a correspondence matrix that reflected label agreement. A reciprocal matrix was constructed by computing, for each predicted label, the distribution of manual annotations to ensure symmetric comparison. Averaging the two matrices yielded a balanced concordance score, providing a robust measure of correspondence between manual annotations and transferred labels assigned to our MERFISH data.

**Correlation between MERFISH experiments**

To assess reproducibility across MERFISH experiments, we computed the mean gene expression profile per experiment and visualized pairwise correlations between replicates as heatmaps. Spearman’s rank correlation coefficients were used to quantify the agreement in gene expression levels across replicates.

**Co-embedding of MERFISH with snRNA-seq**

To directly compare our transcriptomic data from MERFISH with the reference data from snRNA-seq, we jointly embedded both datasets into a shared latent space using the SeuratIntegration class from the ALLCools package (13) for integrated visualization and correspondence analysis. Specifically, both datasets were restricted to a common set of 467 genes shared across MERFISH gene panels to ensure feature compatibility. For each dataset, total counts per cell were normalized, log-transformed, and scaled with the Scanpy standard preprocessing pipeline as described above. Cell clusters annotated with Unassigned and Unresolved in the snRNA-seq dataset were excluded from downstream analysis. To harmonize label granularity, fine-grained snRNA-seq cell type annotations were merged into nine broader categories (Astrocytes, Oligodendrocyte lineage, Ependymal, Endothelial, VLMCs, Microglia, Excitatory neurons, Inhibitory neurons, and Motor neurons). MERFISH subtype annotations were collapsed into the same categories. The preprocessed datasets were concatenated into a joint AnnData object. Dimensionality reduction was performed with PCA, followed by batch correction with Harmony (14).

Integration was performed in the corrected PC space using canonical correlation analysis (CCA), with integration anchors defined as mutual nearest neighbors across datasets. To address scalability constraints, CCA was computed on a random subset of MERFISH cells to match in size to the snRNA-seq dataset, before being applied to all MERFISH cells. The integrated low-dimensional embeddings were used to construct a shared KNN graph, which was subsequently used to compute a two-dimensional UMAP for direct visualization of transcriptomic organization and label concordance between the MERFISH cell clusters and snRNA-seq clusters.

**Relative depth analysis**

To ensure accurate depth estimation, only intact sections free of tissue tears were included in the spatial analyses. Spatial coordinates of individual cells were obtained from the processed MERFISH dataset. Raw depth of a cell was defined as the vertical distance between its centroid and the dorsal surface of the section. The dorsal boundary was defined manually for each section using 7–14 anchor points placed along the visible dorsal contour. To construct a continuous surface estimate, anchor points were fit with cubic spline interpolation. To standardize orientation across sections, an anchor-based rotation procedure was implemented. Specifically, the left-most and right-most anchor points were identified, and their centroids were used to compute the angle between the anchor-defined dorsal edge and the original horizontal axis. A rotation matrix was then applied to re-align the section such that the dorsal surface was horizontal. Before rotation, all cell coordinates were translated so that the midpoint of the two anchor points was set as the new origin. Cell depth was computed relative to the fitted dorsal surface. For each cell centroid, the spline-estimated dorsal y-coordinate at its x-position was subtracted from the cell's y-coordinate to obtain a raw depth (μm). Relative depth was defined as:

$$Relative\ depth = \frac{d_i}{d_{max}}$$

Where $d_i$ is the raw depth of a given cell $i$, and $d_{max}$ is the maximum depth of all cells within that section. This normalization ensured comparability across sections of varying shapes. Boundary fitting, spline interpolation, and depth calculation were performed using custom functions (fit_northern_boundary, anchor_based_rotation, compute_depths). For quality control, diagnostic plots, including spline fits of the dorsal boundary, depth-colored scatter maps of cells within each section, and violin plots of cell-type depth distributions were also generated. Relative depths of all neuronal subtypes were averaged across sections and plotted for visualization.

**Cell-cell proximity analysis**

To identify statistically enriched or depleted spatial co-localization between MERFISH cell types, spatial proximity analysis was performed by computing pairwise Euclidean distances between all cell soma centroid coordinates within all intact sections. For each cell, its neighboring cells were defined as all cells located within a 60 μm radius. The composition of neighboring cell types of each cell was aggregated across all cell types to construct an observed cell–cell proximity matrix, where each entry represented the frequency with which one cell type was proximal to another cell type. To estimate a null expectation for cell proximity independent of spatial arrangement, we performed permutation testing by randomly shuffling cell type labels across all cells while preserving their spatial positions. This procedure was repeated for 100 iterations to compute the average expected proximity matrix. Observed proximity frequencies were normalized to expected frequencies using a $\log 2$ ratio, clipped to the range [-2, 2] for visualization. Statistical significance of observed versus expected interactions was assessed using Wilcoxon rank-sum tests comparing neighbor counts from observed versus permuted datasets. Resulting p-values were adjusted for multiple hypothesis testing with the Benjamini-Hochberg procedure. False discovery rate (FDR)-adjusted p-values were capped at 4 for better visualization. Section-level results were combined by averaging normalized proximity matrices and FDR scores across all sections analyzed.

**Cell-label transfer for the MERFISH pain dataset**

To annotate cell types in the neuropathic pain dataset, we implemented an ensemble label transfer strategy that integrates four state-of-the-art methods: SingleR, Tangram (15), Seurat, and Robust Cell Type Decomposition (RCTD) (16). Each method independently predicted labels for queried cells in the pain dataset using annotations from our manually annotated MERFISH dataset (referred to as the reference dataset below).

Tangram: Both reference and pain datasets were normalized by total counts (scanpy.pp.normalize_total) and log-transformed (scanpy.pp.log1p) using the Scanpy standard preprocessing pipeline as described above. We subsampled 50,000 cells from the reference dataset prior to training, as Tangram requires a tractable reference size to avoid GPU memory overflow. Preprocessing and gene-space alignment were performed with tg.pp_adatas, which harmonizes gene features across the reference and pain datasets. Cell-to-space mapping was performed with tg.map_cells_to_space using a uniform density prior to evenly distributing reference cells across spatial coordinates and leveraging GPU acceleration for efficient optimization. The resulting mapping object was projected back onto the pain dataset using tg.project_cell_annotations, thereby transferring cell type annotations from the reference to the pain dataset.

SingleR: Reference and pain datasets were first converted into Seurat objects normalized with LogNormalize, and converted into SingleCellExperiment objects for compatibility with SingleR. For the reference dataset, the MERFISH cell type annotations available in the metadata were used as ground-truth labels. SingleR was then applied with the Wilcoxon test (de.method = 'wilcox') to identify discriminative genes, using the top 10 markers per comparison (de.n = 10). Expression profiles of cells in the pain dataset were compared against those in the reference dataset, and the most correlated reference annotation was assigned to the individual cell in the pain dataset based on the computed similarity scores.

Seurat: Both datasets were converted into Seurat objects. Expression matrices were transposed to match the expected genes-by-cells format and stored as integer matrices. Each dataset was independently processed through standard Seurat workflows, including normalization (NormalizeData), identification of variable features (FindVariableFeatures), scaling (ScaleData), and dimensionality reduction via PCA. Cross-dataset correspondences were identified using FindTransferAnchors with 30 PCs, and cell type labels from the reference were then transferred to the pain dataset using TransferData.

RCTD: The reference count matrix and metadata were first transposed to ensure the genes-by-cells format and stored as an integer matrix. Metadata containing MERFISH cell type annotations were parsed to construct factorized labels, and cells with less than 20 UMIs were excluded prior to constructing the reference object. For the pain dataset, raw counts were similarly processed into an integer matrix. Spatial coordinates were extracted and combined with the count matrix to generate a SpatialRNA object. With these objects, an RCTD model was initialized with create.RCTD specifying a minimum UMI threshold of 10. Deconvolution was performed in doublet mode using run.RCTD to account for potential mixed-cell signals in MERFISH pixels. Predicted cell type weights were then extracted for integration into the ensemble label transfer framework.

Majority voting: To enhance the overall accuracy of cell type annotation, we employed a weighted majority voting framework that integrates predictions from multiple label transfer methods. Weights for each method were optimized using a 70/30 training–testing split of the reference dataset. During training, candidate weight combinations were evaluated based on their ability to maximize label prediction accuracy on the testing set. Incorporation into the weighted majority voting scheme improved performance by approximately 5%, reaching an overall accuracy of 78.4%. The optimized ensemble model was subsequently applied to the neuropathic pain dataset for automatic cell type annotation.

**Cell-cell communication (CCC)**

To investigate intercellular communication mediated by ligand–receptor (L-R) interactions, we applied NICHES (17) [to] spinal cord sections from a single MERFISH experiment. Raw counts were normalized with SCTransform in Seurat. To enhance weak biological signals and reduce clustering artifacts related to gene sampling, data imputation was performed with RunALRA from SeuratWrappers. To mitigate data inflation, structural downsampling for each cell type was performed within NICHES. Cell–cell signaling was computed using the RunNICHES function in CellToCell mode. All L-R mechanisms were obtained from the Omnipath database via OmnipathR::import_ligrecextra_interactions function in NICHES, restricted to experimentally validated interactions. After computing L-R expression values, an L-R assay was created with rows as the L-R mechanism and columns as sender-receiver cell-cell pairs. L-R mechanisms were normalized across cell-cell pairs using the Seurat ScaleData function, with expression in at least 25% of cell–cell pairs retained. Niche-specific signaling interactions were identified using Seurat FindAllMarkers with the ROC test applied to individual cell type-cell type interaction groups. For each group, the top five L-R mechanisms ranked by AUC were selected for differential expression analysis. To prioritize L-R markers, we ranked L-R mechanisms by a defined composite score, derived from the FindAllMarkers output as:

$$S = (p_{in} - p_{out}) \times AUC \times \overline{\log_2 FC}$$

Where $p_{in}$ and $p_{out}$ denote the proportions of the given sender-receiver pair expressing the L-R interaction within and outside the target group, $AUC$ denotes the area under the ROC curve obtained from the FindAllMarkers function (test.use = "roc"), and $\overline{\log_2 FC}$ is the average $\log 2$ fold change in L-R interaction expression between the target group and the remaining cell-cell pairs. DoHeatmap function was used to visualize the expression of the top L-R mechanisms.

Neuron subtypes were assigned to respective lamina based on prior knowledge as, lamina I: DH-ex-Reln/Nmur2, DH-ex-Reln/Npff, DH-ex-Gpr83, DH-in-Pdyn/Gal, DH-in-Pdyn, DH-ex-Grpr; lamina II: DH-ex-Sox5, DH-ex-Reln, DH-ex-Nmu/Tac2, DH-ex-Grp, DH-ex-Sox5/Tac1, DH-in-Rorb, DH-ex-Grpr; lamina III: DH-ex-Cpne4, DH-ex-Tac2, DH-ex-Prkcg/Cck, DH-ex-Prkcg/Nts, DH-ex-Prkcg/Rxfp1, DH-ex-Rreb1, DH-ex-Maf/Cck, DH-ex-Maf/Slc17a8, DH-ex-Maf/Cpne4, DH-in-Rorb, DH-in-Npy, DH-in-Npy2r, DH-in-Kcnip2, DH-in-Cdh3, DH-in-Klhl14.

**Generation and assessment of neuropathic pain**

We produced spared nerve injury (SNI) mice as models for neuropathic pain, and sham-surgery mice as the control group. In both cases, 8-10-week-old male and female mice were anesthetized using isoflurane. The hair on the left hind leg was removed by Nair, and the exposed skin was disinfected with 70% ethanol and then Betadine. After an incision was made above the knee joint, muscles were separated to expose the sciatic nerve bundle. For SNI mice, a transection was made between two knots, where one tied the tibial and common peroneal nerves using non-absorbable silk suture without touching the sural nerve, and the other was tied at a lower position with the absorbable Vicryl suture. For sham mice, no operation was performed on the nerves. The muscles were then pulled together, and the skin was stitched. A liquid bandage was applied to the skin.

The dynamic brush test was used to assess the development of neuropathic pain. Mice were habituated for 30 min on a metal grid in a clear acrylic chamber for 2 days. Three gentle brushes were applied to the lateral part of the left hind paw using a paintbrush. Brushing continued with 5-10 s intervals for 3 min per trial. Three trials were performed for each animal. The responses were graded with a score from 0 to 3: 0 for no response, 1 for sustained lifting or withdrawal of the ipsilateral hindpaw, 2 for lateral kicking, and 3 for licking. The maximal response for each trial was considered the score for that trial, and the scores for all three trials were averaged. Sham and SNI mice were perfused without sensory stimulation, and their spinal cords were sectioned for MERFISH imaging.

**Perturbation Analysis Using Augur**

To identify the neuronal subtypes most transcriptionally perturbed under neuropathic pain, we applied Augur, a machine learning–based framework for quantifying cell-type-specific sensitivity to experimental perturbations. Augur uses a subsampling strategy to train a random forest classifier that predicts sample identity from transcriptional profiles. The area under the receiver operating characteristic curve (AUC) was aggregated across all neuronal subtypes to generate a perturbation ranking, where higher AUC values indicate that a given subtype has more separability between sham and SNI conditions. Augur was used as a prioritization framework to rank relative cell-type responsiveness to perturbation, with running parameters: n_subsamples = 50, subsample_size = 15, feature_perc = 0.5. In this framework, an AUC of 0.5 corresponds to chance-level separability, whereas a higher AUC indicates that cells from the two conditions are more separable in the feature space and therefore that the cell type shows a stronger molecular response to the perturbation. Neuronal subtypes that contain fewer than 15 cells in either condition were discarded to ensure the robustness of the results.

### Neuronal activity between SNI and sham mice

The expression of activity-related genes (ARG), defined as the ARG score, was measured to assess neuronal activity in SNI and sham mice across spinal cord neuron subtypes. Cells were subsetted to include only neurons from the left side of the lumbar sections, the regions of interest. The ARG score was calculated per cell as the mean z-scored expression of 6 ARGs (*Arc*, *Junb*, *Fos*, *Fosb*, *Npas4*, *Nr4a1*), thereby standardizing for baseline expression differences. To control for section-level variability, ARG scores were averaged per section for pseudobulk analysis. Wilcoxon rank-sum tests were performed to compare the ARG score of each neuronal subtype between SNI and sham conditions.

### CellChat analysis between SNI and sham mice

To characterize and compare intercellular communication networks underlying control and pain conditions, we applied CellChat, a computational framework that infers cell–cell signaling networks from transcriptomic profiles using curated ligand–receptor interactions. CellChat leverages CellChatDB, a manually curated database of literature-supported ligand–receptor pairs to predict potential communication links between cell groups based on expression of corresponding ligands and receptors. For each condition, we initialized a CellChat object using the spatial transcriptomic expression matrix and associated cell type annotations. Raw gene expression was normalized with the normalizeData function from CellChat, which performs library-size scaling and log transformation prior to downstream analysis. When computing communication probabilities with computeCommunProb, we set trim = 0.1 implying that the average gene expression was zero if the percentage of expressing cells in one group was less than 10%. For our MERFISH data, we used conversion.factor = 1 since the spatial coordinates are already in micrometers and set contact.range = 50 to restrict the contact-dependent signaling. Following inference of cell–cell communication networks, significant interactions were retained through CellChat's internal statistical procedures, which filter out non-significant links. Communication networks at the signaling-pathway level were derived by aggregating the communication probabilities of all ligand-receptor pairs associated with each pathway between cell type pairs. Based on the interaction strength of each signaling pathway, we used rankNet from CellChat to compare the overall information flow of each signaling pathway and identify altered signaling pathways with distinct interaction strengths between conditions. Finally, we visualized the communication probabilities, which are assigned as edge weights to quantify the interaction strength for each condition in circle plots.

### Quantification and statistical analysis

Statistical tests were described in the respective figure legends and Methods sections. Welch's t-tests were used for differences in the percentage of cell types between males and females. Wilcoxon rank-sum tests were used for the difference in ARG scores between sham and SNI groups. Wilcoxon rank-sum tests with FDR-adjusted p-values were used for proximity analysis. Group differences were considered statistically significant at $p < 0.05$. Significance levels are indicated as follows: $*p < 0.05$; $**p < 0.01$; $***p < 0.001$; n.s., not significant. All data are presented as the mean ± the standard error of the mean (SEM), unless otherwise noted in the figure legends. Statistical analyses were performed using GraphPad Prism software and R.

**Interactive web-based browser for MERFISH data**

To create a web browser of our MERFISH data, we used the WebAtlas pipeline (18), which leverages Vitessce (19) to visualize spatial transcriptomic data in the Zarr format. Data were uploaded to Github for easy access and retrieval. WebAtlas retrieves the data using its web portal hosted by the Wellcome Sanger Institute, and provides interactive visualizations including UMAP plots, cell-type annotation, and gene-expression distributions. All of the visualizations can be found at: https://github.com/shanmeltzer/Spinal_MERFISH.

**Supplemental Figures**

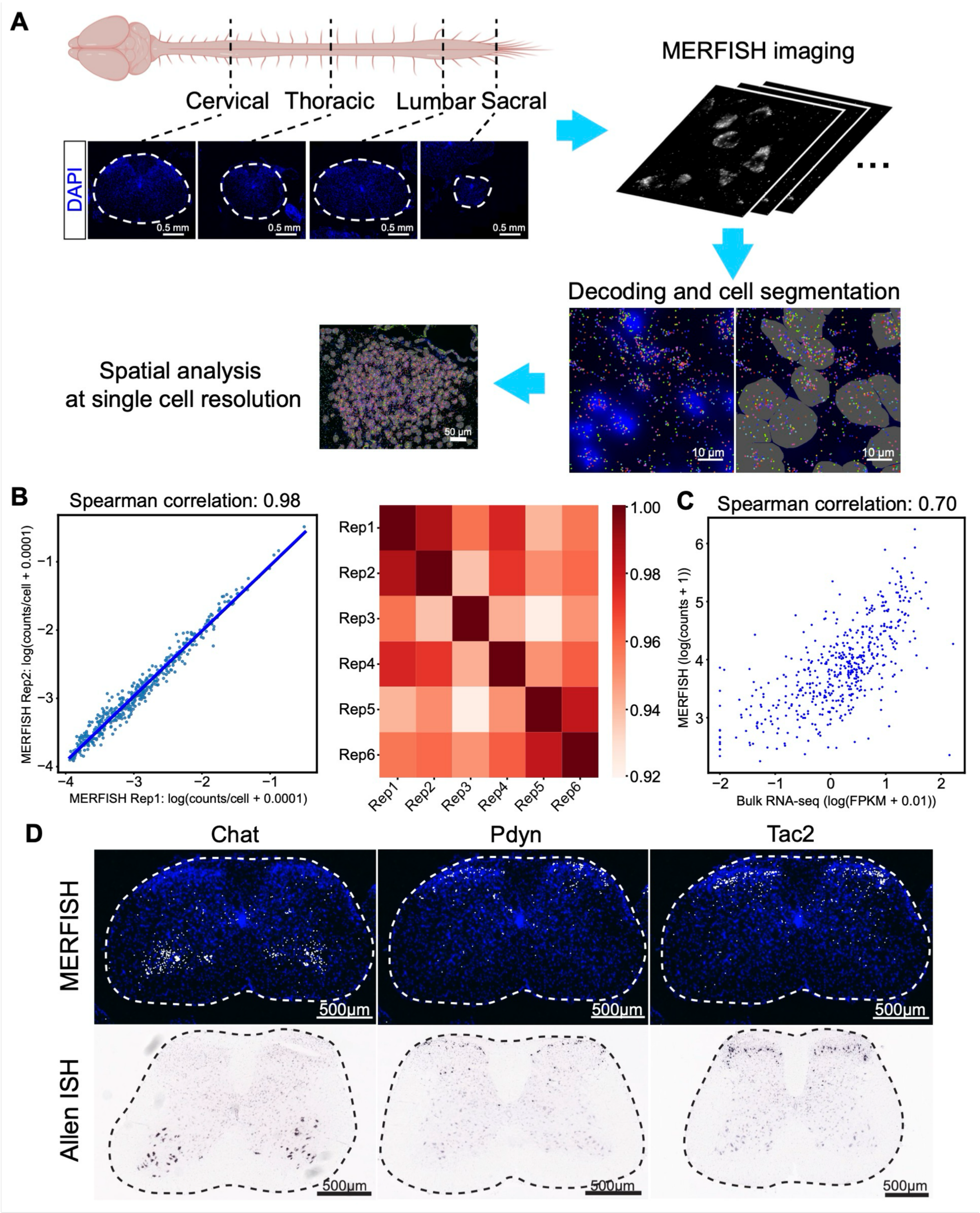


**Figure S1. Workflow and quality control of MERFISH profiling of the mouse spinal cord. Related to Figure 1.**

(A) Workflow of MERFISH profiling of the mouse spinal cord, including MERFISH imaging, decoding, cell segmentation, and data analysis. Sections of cervical, thoracic, lumbar, and sacral levels were collected and imaged for each animal. (B) Scatterplot of the Spearman correlation coefficient between average RNA counts per cell of individual genes measured with MERFISH in independent experiments. Heatmap of Spearman correlation coefficients for all MERFISH experiments (n = 6 technical replicates), indicating strong correlation between each MERFISH measurement. (C) Scatterplot of the Spearman correlation coefficient between RNA counts measured by

MERFISH and average expression determined from bulk RNA-sequencing of the mouse spinal cord. (D) Spatial gene expression of three representative genes detected by MERFISH is shown on the top. In situ hybridization (ISH) data from the Allen Spinal Cord Reference Atlas are shown at the bottom. Scale bars, 500 µm.

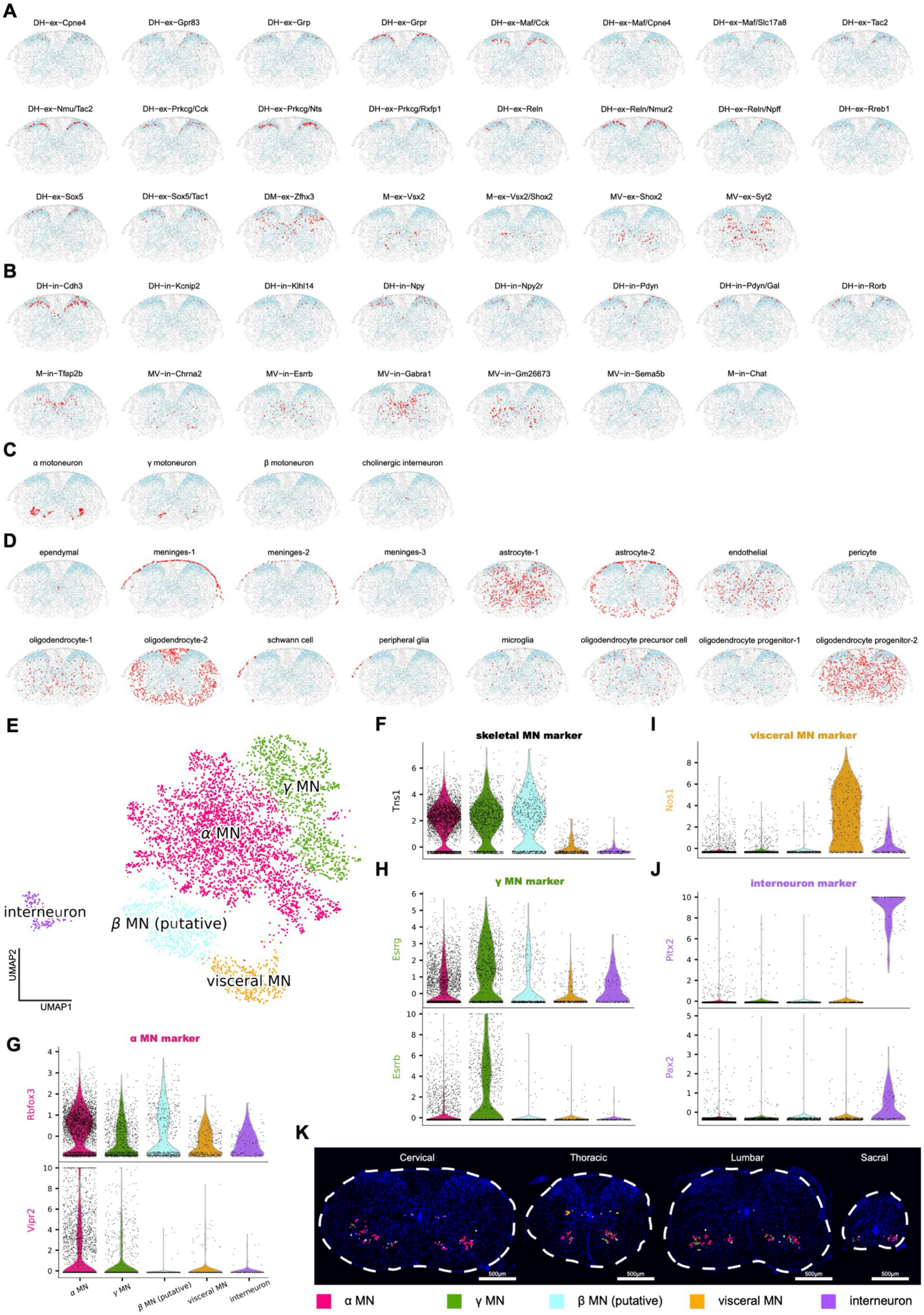

**Figure S2. Spatial locations of all MERFISH cell types and characterization of spinal cord cholinergic neurons. Related to Figure 1.**
(A-D) Excitatory (A), inhibitory (B), cholinergic (C), and non-neuronal types (D) on the representative cervical section. Red dots mark the indicated cell subtypes, light blue dots mark neurons, and gray dots mark non-neuronal cells. No M-ex-Neurod2 and visceral motoneuron subtypes were identified in this section. (E) UMAP visualization of identified cholinergic neuron subtypes by MERFISH. (F-J) Violin plots showing marker gene distributions for skeletal motoneurons (F), α motoneurons (G), γ motoneurons (H), visceral motoneurons (I), and cholinergic interneurons (J). (K) Spatial distributions of cholinergic neuron subtypes at different axial levels.

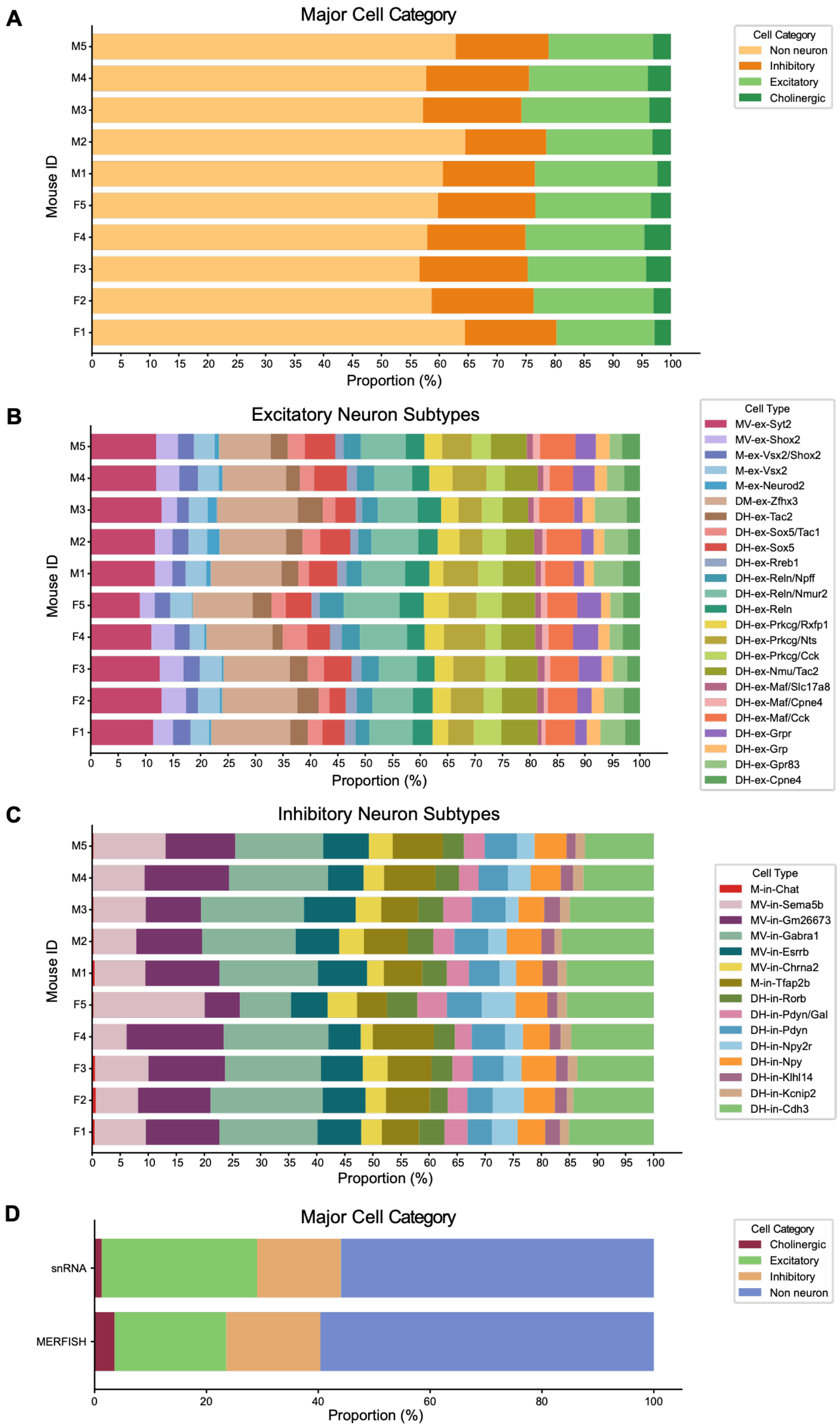


**Figure S3. Compositions of cell categories and neuron subtypes. Related to Figure 1.**
(A) Stacked bar plots of average proportions of major cell categories across 10 animals in the MERFISH dataset. (B-C) Stacked bar plots of average proportions of excitatory (B) and inhibitory (C) neuron subtypes across animals in the MERFISH dataset. (D) Stacked bar plots of average proportions of major cell categories in MERFISH and snRNA-seq datasets.

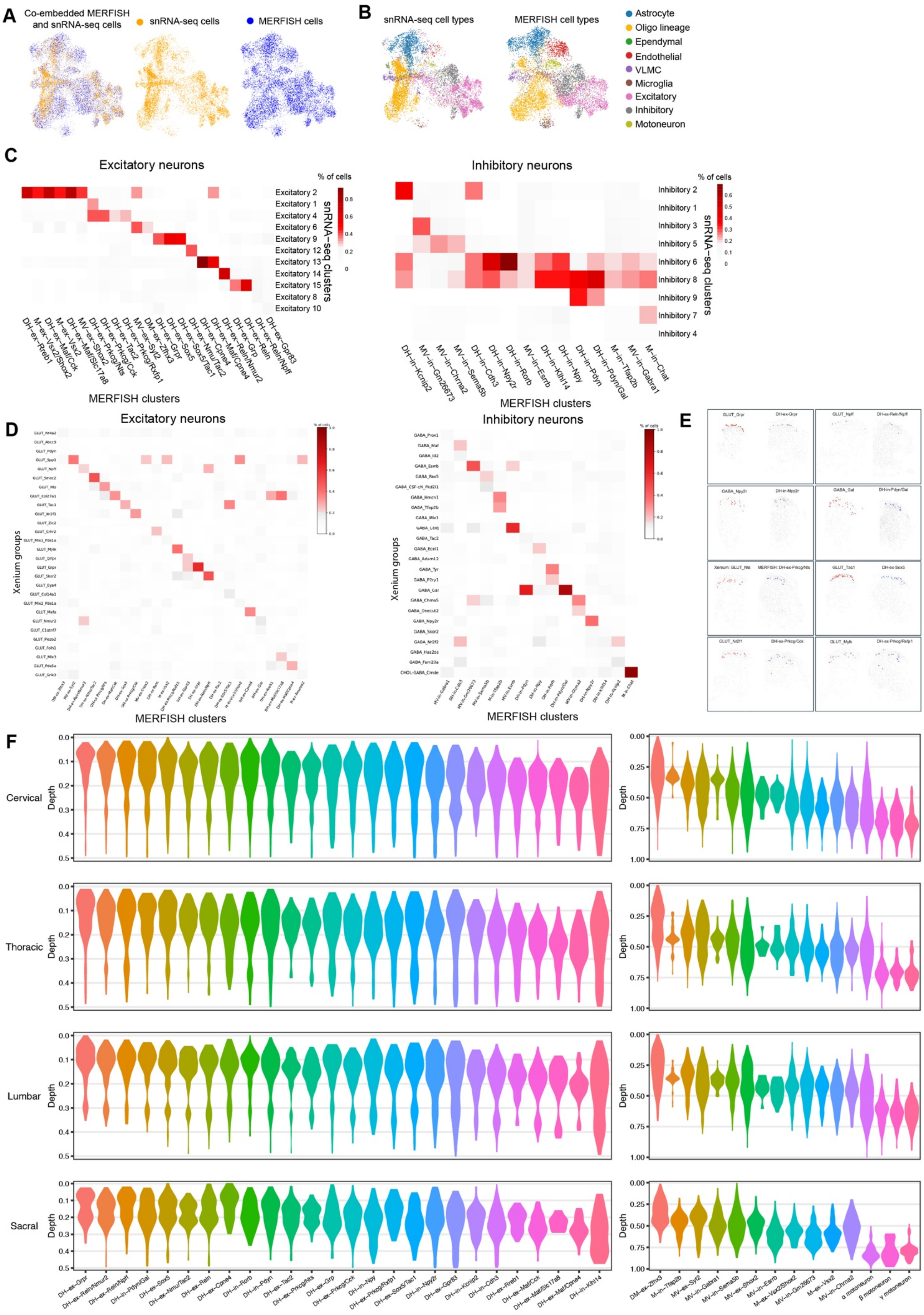

A
Co-embedded MERFISH and snRNA-seq cells
snRNA-seq cells
MERFISH cells
B
snRNA-seq cell types
MERFISH cell types
Astrocyte
Oligo lineage
Ependymal
Endothelial
VLMC
Microglia
Excitatory
Inhibitory
Motoneuron
C
Excitatory neurons
% of cells
Excitatory 2
Excitatory 1
Excitatory 4
Excitatory 6
Excitatory 9
Excitatory 12
Excitatory 13
Excitatory 14
Excitatory 15
Excitatory 8
Excitatory 10
snRNA-seq clusters
DH-ex-Rreb1
M-ex-Vsx2/Shox2
M-ex-Vsx2
DH-ex-Maf/Cck
DH-ex-Maf/Slc17a8
MV-ex-Shox2
DH-ex-Prkcg/Nts
DH-ex-Prkcg/Cck
DH-ex-Prkcg/Rxfp1
DH-ex-Tac2
MV-ex-Syt2
DM-ex-Zfhx3
DH-ex-Grpr
DH-ex-Sox5
DH-ex-Sox5/Tac1
DH-ex-Nmu/Tac2
DH-ex-Cpne4
DH-ex-Maf/Cpne4
DH-ex-Reln/Nmur2
DH-ex-Grp
DH-ex-Reln
DH-ex-Reln/Npff
DH-ex-Gpr83
MERFISH clusters
Inhibitory neurons
% of cells
Inhibitory 2
Inhibitory 1
Inhibitory 3
Inhibitory 5
Inhibitory 6
Inhibitory 8
Inhibitory 9
Inhibitory 7
Inhibitory 4
snRNA-seq clusters
DH-in-Kcnip2
MV-in-Gm26673
MV-in-Chrna2
MV-in-Sema5b
DH-in-Cdh3
DH-in-Npy2r
DH-in-Rorb
MV-in-Esrrb
DH-in-Klhl14
DH-in-Npy
DH-in-Pdyn
DH-in-Pdyn/Gal
M-in-Tfap2b
MV-in-Gabra1
M-in-Chat
MERFISH clusters
D
Excitatory neurons
Xenium groups
MERFISH clusters
Inhibitory neurons
Xenium groups
MERFISH clusters
E
F
Cervical
Thoracic
Lumbar
Sacral
Depth
DH-ex-Grpr
DH-ex-Reln/Nmur2
DH-ex-Reln/Npff
DH-in-Pdyn/Gal
DH-ex-Sox5
DH-ex-Nmu/Tac2
DH-ex-Reln
DH-ex-Cpne4
DH-in-Rorb
DH-in-Pdyn
DH-ex-Tac2
DH-ex-Prkcg/Nts
DH-ex-Grp
DH-ex-Prkcg/Cck
DH-in-Npy
DH-ex-Prkcg/Rxfp1
DH-ex-Sox5/Tac1
DH-in-Npy2r
DH-ex-Gpr83
DH-in-Kcnip2
DH-in-Cdh3
DH-ex-Rreb1
DH-ex-Maf/Cck
DH-ex-Maf/Slc17a8
DH-ex-Maf/Cpne4
DH-in-Klhl14
DM-ex-Zfhx3
M-in-Tfap2b
MV-ex-Syt2
MV-in-Gabra1
MV-in-Sema5b
MV-ex-Shox2
MV-in-Esrrb
M-ex-Vsx2/Shox2
MV-in-Gm26673
M-ex-Vsx2
MV-in-Chrna2
α motoneuron
β motoneuron
γ motoneuron

**Figure S4. MERFISH and snRNA-seq clusters of spinal cord cells are consistent. Related to Figure 1.**

(A) UMAP visualizations of co-embedded MERFISH and snRNA-seq (Russ et al., 2021) with a randomly down-sampled subset of MERFISH cells matching the cell number of the snRNA-seq dataset. UMAP of superimposed (left), only snRNA-seq (middle), and only MERFISH cells (right) are shown separately. (B) UMAP visualizations of major cell classes determined with snRNA-seq (left) and MERFISH (right), showing correspondence of cell classes between MERFISH and snRNA-seq datasets. (C and D) Heatmap of the percentage of MERFISH excitatory and inhibitory neuron subtypes (columns) assigned to snRNA-seq neurons (rows), showing the correspondence of identified cell types between MERFISH and snRNA-seq (C) datasets and between MERFISH and Xenium (D) datasets. (E) Spatial distribution of representative groups in the MERFISH and corresponding Xenium datasets, showing the similar spatial distribution of neuronal types between the two datasets. (F) Violin plots showing distributions of the relative depth of respective neuron subtypes across axial levels. The maximum distance is normalized to 1 for each section.

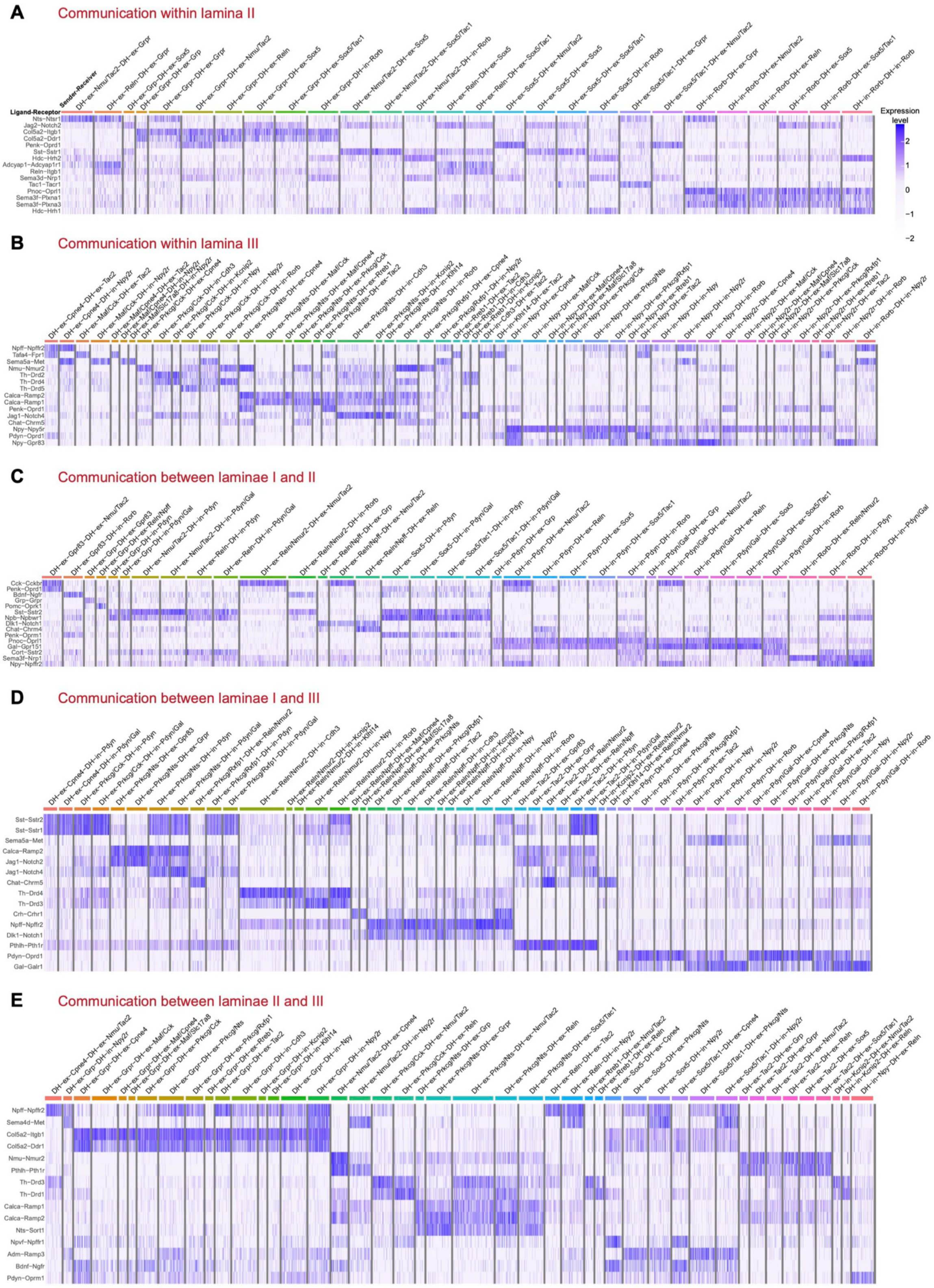

A
Communication within lamina II
Sender-Receiver
Ligand-Receptor
Nts-Ntsr1
Jag2-Notch2
Col5a2-Itgb1
Col5a2-Ddr1
Penk-Oprd1
Sst-Sstr1
Hdc-Hrh2
Adcyap1-Adcyap1r1
Reln-Itgb1
Sema3d-Nrp1
Tac1-Tacr1
Pnoc-Oprl1
Sema3f-Plxna1
Sema3f-Plxna3
Hdc-Hrh1
Expression level
B
Communication within lamina III
Npff-Npffr2
Tafa4-Fpr1
Sema5a-Met
Nmu-Nmur2
Th-Drd2
Th-Drd4
Th-Drd5
Calca-Ramp2
Calca-Ramp1
Penk-Oprd1
Jag1-Notch4
Chat-Chrm5
Npy-Npy5r
Pdyn-Oprd1
Npy-Gpr83
C
Communication between laminae I and II
Cck-Cckbr
Penk-Oprd1
Bdnf-Ngfr
Grp-Grpr
Pomc-Oprk1
Sst-Sstr2
Npb-Npbwr1
Dlk1-Notch1
Chat-Chrm4
Penk-Oprm1
Pnoc-Oprl1
Gal-Gpr151
Cort-Sstr2
Sema3f-Nrp1
Npy-Npffr2
D
Communication between laminae I and III
Sst-Sstr2
Sst-Sstr1
Sema5a-Met
Calca-Ramp2
Jag1-Notch2
Jag1-Notch4
Chat-Chrm5
Th-Drd4
Th-Drd3
Crh-Crhr1
Npff-Npffr2
Dlk1-Notch1
Pthlh-Pth1r
Pdyn-Oprd1
Gal-Galr1
E
Communication between laminae II and III
Npff-Npffr2
Sema4d-Met
Col5a2-Itgb1
Col5a2-Ddr1
Nmu-Nmur2
Pthlh-Pth1r
Th-Drd3
Th-Drd1
Calca-Ramp1
Calca-Ramp2
Nts-Sort1
Npvf-Npffr1
Adm-Ramp3
Bdnf-Ngfr
Pdyn-Oprm1

**Figure S5. Cell-cell communication of neuron populations through ligand-receptor interactions within and between laminae of the spinal cord dorsal horn. Related to Figure 5.**

(A-E) Heatmaps showing the expression of the next top 15 ligand–receptor interactions identified among neuronal subtypes located within lamina II (A), lamina III (B), and between laminae I and II (C), laminae I and III (D), laminae II and III (E). Rows represent ligand–receptor pairs, and columns correspond to sender–receiver neuron populations. The color scale indicates normalized expression levels of the ligand-receptor pair.

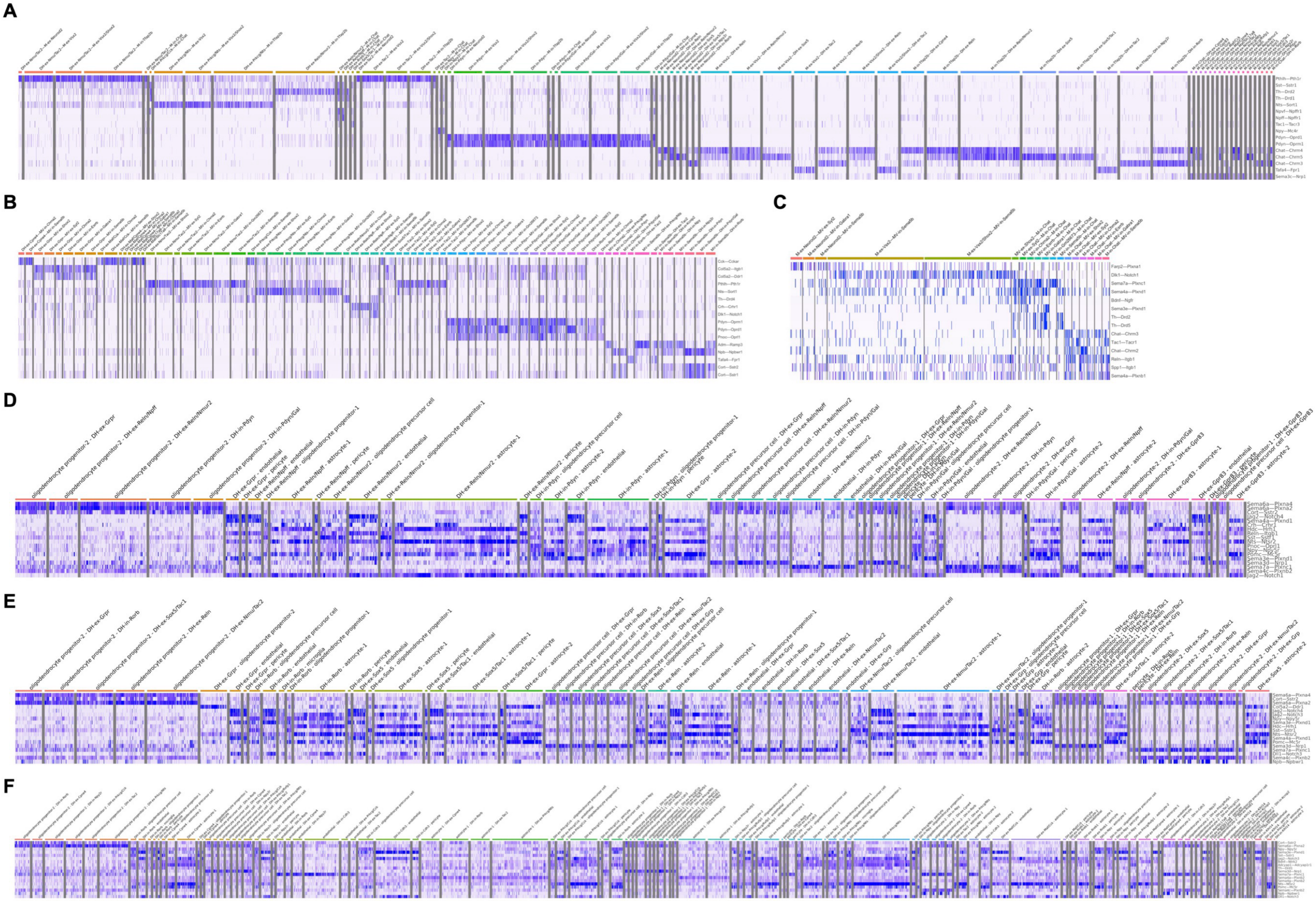
A
B
C
D
E
F

**Figure S6. Cell-cell communication between neuron and non-neuron populations. Related to Figure 5.**

(A-F) Heatmaps showing the expression of ligand-receptor interactions of neuron populations within dorsal and intermediate (A), dorsal and ventral (B), and intermediate and ventral regions (C) of the spinal cord, respectively, and interactions identified among neuron and non-neuron types located in lamina I (D), II (E), and III (F), respectively. Rows represent ligand-receptor pairs, and columns correspond to sender-receiver neuron populations. The color scale indicates normalized expression levels of the ligand-receptor pair.

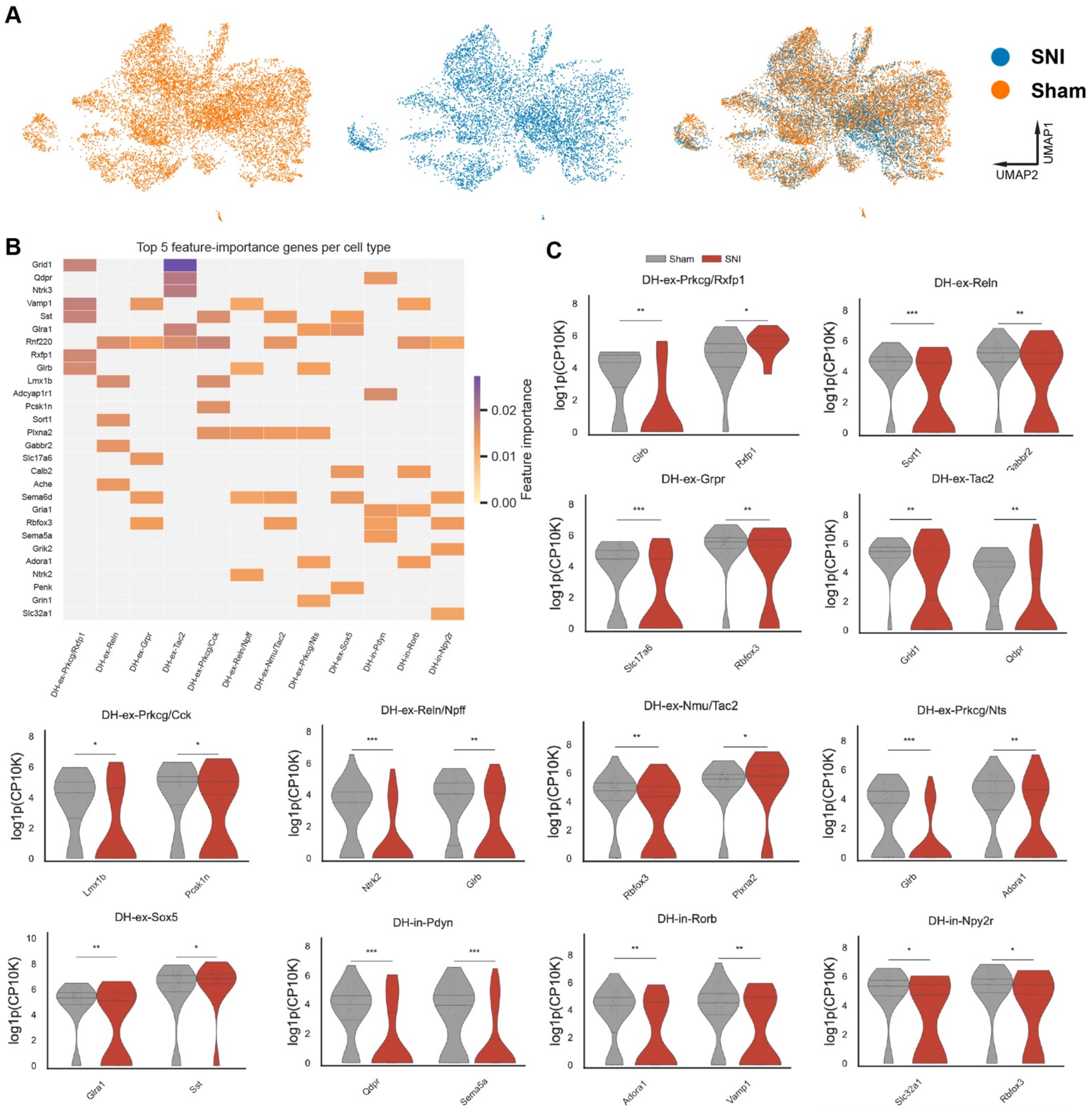


**Figure S7. Transcriptomic changes in sham and SNI spinal cords. Related to Figure 6.**

(A) UMAP visualizations of cells from sham animals (left), SNI models (middle), and from both groups combined (right), showing correspondence between sham and SNI groups. (B) Feature importance genes of Augur cell type prioritizations for left lumbar spinal cord neurons in sham and SNI conditions. (C) Violin plots showing expression levels of the top 2 feature-importance genes for each neuronal subtype in sham and SNI conditions. Wilcoxon rank-sum test, *$p < 0.05$, **$p < 0.01$, ***$p < 0.001$.

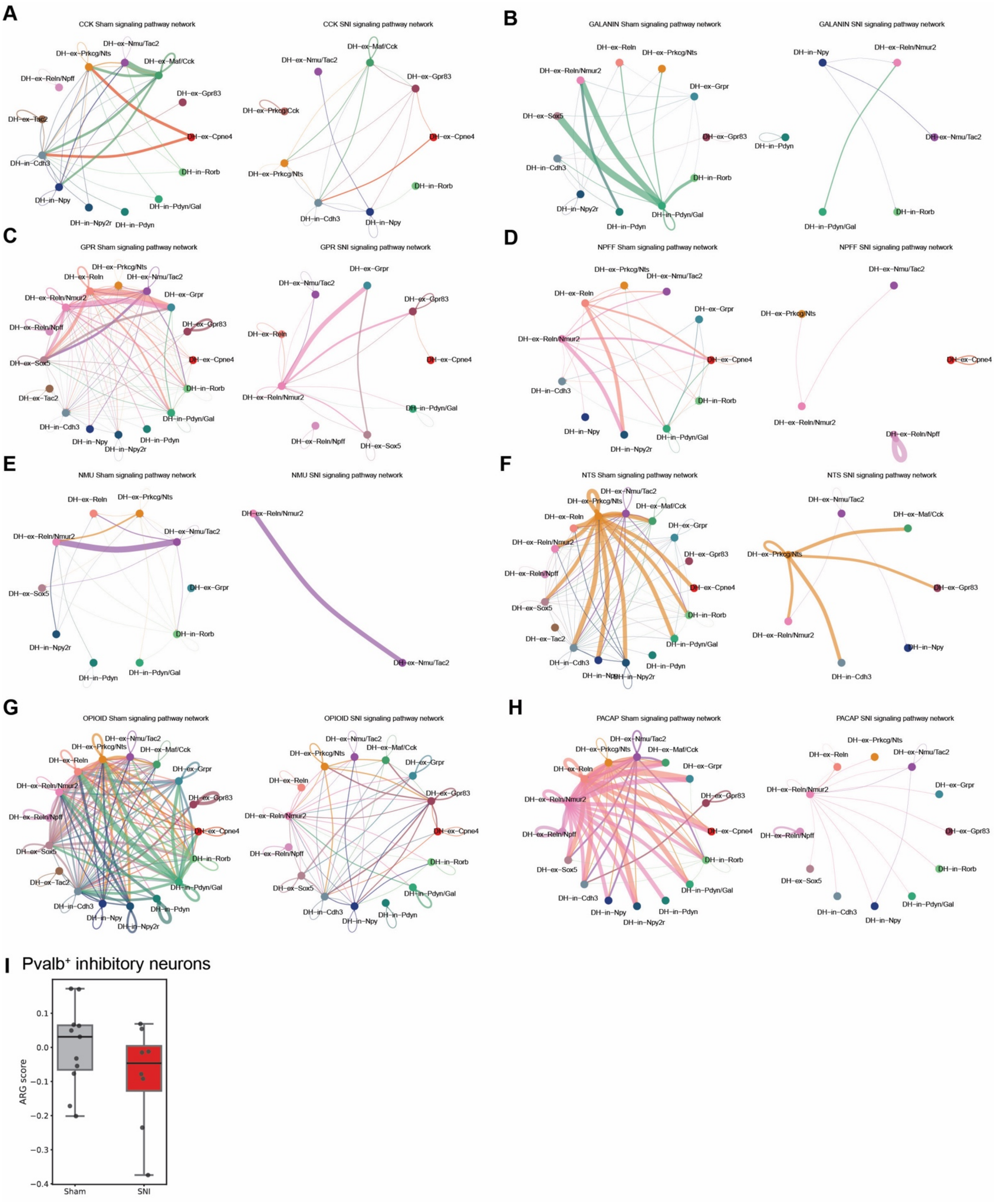


**Figure S8. Communication between neuropeptide signaling in sham and SNI spinal cords. Related to Figure 6.**

(A-H) Detected possible interactions in the neuropeptide signaling pathways between pairs of dorsal horn neuronal subtypes. Edge width represents the communication probability, and each edge is colored to match its sender neuronal subtype, indicating the direction of signal origin. (I) Box plots of ARG scores in Parvalbumin-positive neurons in sham and SNI animals. Not significant determined by Wilcoxon rank-sum tests.

Tables S1 MERFISH Gene Panel

| Gene name | Category |
|---|---|
| Abcc9 | Pericytes |
| Ache | neurotransmitter-Choline |
| Adamts2 | DH Neurons |
| Adamts5 | DH Neurons |
| Adcyap1 | NP-GPCR/NPR genes |
| Adcyap1r1 | NP-GPCR/NPR genes |
| Adm | Ligand-receptor pairs-CGRP |
| Adm2 | Ligand-receptor pairs-CGRP |
| Adora1 | neurotransmitter-adenosin |
| Adora2a | neurotransmitter-adenosin |
| Adora2b | neurotransmitter-adenosin |
| Adora3 | neurotransmitter-adenosin |
| Adra1a | neurotransmitter-Norepinephrine and epinephrine |
| Adra1b | neurotransmitter-Norepinephrine and epinephrine |
| Adra1d | neurotransmitter-Norepinephrine and epinephrine |
| Adra2a | neurotransmitter-Norepinephrine and epinephrine |
| Adra2b | neurotransmitter-Norepinephrine and epinephrine |
| Adra2c | neurotransmitter-Norepinephrine and epinephrine |
| Adrb1 | neurotransmitter-Norepinephrine and epinephrine |
| Adrb2 | neurotransmitter-Norepinephrine and epinephrine |
| Adrb3 | neurotransmitter-Norepinephrine and epinephrine |
| Agt | Neuroactive Peptides-Angiotensin |
| Agtr1a | Neuroactive Peptides-Angiotensin |
| Agtr1b | Neuroactive Peptides-Angiotensin |
| Agtr2 | Neuroactive Peptides-Angiotensin |
| Aif1 | Microglia |
| Aldh7a1 | neurotransmitter-histamine |
| Aoc1 | neurotransmitter-histamine |
| Apela | Neuroactive Peptides-Apelin |
| Apln | Neuroactive Peptides-Apelin |
| Aplnr | Neuroactive Peptides-Apelin |
| Arc | Activity relavant genes |
| Bdkrb1 | Neuroactive Peptides-Bradykinin |
| Bdkrb2 | Neuroactive Peptides-Bradykinin |
| Bdnf | Ligand-receptor pairs-neurotrophin |
| Bicc1 | Meninges-1 |
| Bnc2 | DH Neurons |
| Brs3 | Neuroactive Peptides |
| C1ql1 | OPC |
| C3 | Neuroactive Peptides-Anaphylatoxin and TLQP-21 |
| C3ar1 | Neuroactive Peptides-Anaphylatoxin and TLQP-21 |
| C4a | Neuroactive Peptides-Anaphylatoxin and TLQP-21 |

| | |
|---|---|
| C4b | Neuroactive Peptides-Anaphylatoxin and TLQP-21 |
| C5ar1 | Neuroactive Peptides-Anaphylatoxin and TLQP-21 |
| C5ar2 | Neuroactive Peptides-Anaphylatoxin and TLQP-21 |
| Calb1 | DH Neurons |
| Calb2 | DH Neurons |
| Calca | Ligand-receptor pairs-CGRP |
| Calcb | Ligand-receptor pairs-CGRP |
| Calcr | Ligand-receptor pairs-CGRP |
| Calcrl | Ligand-receptor pairs-CGRP |
| Car12 | DH Neurons |
| Ccbe1 | DH Neurons |
| Cck | DH Neurons+NP-GPCR/NPR genes |
| Cckar | NP-GPCR/NPR genes |
| Cckbr | NP-GPCR/NPR genes |
| Ccn2 | Ligand-receptor pairs-neurotrophin |
| Ccn3 | Ligand-receptor pairs-Semaphorin |
| Cd72 | Ligand-receptor pairs-Semaphorin |
| Cdh19 | Peripheral glia |
| Cdh3 | DH Neurons |
| Cep112 | DH Neurons |
| Cfap43 | Ependymal |
| Chat | preganglionic cells (PGC)+DH Neurons |
| Chrm1 | neurotransmitter-Choline |
| Chrm2 | neurotransmitter-Choline |
| Chrm3 | neurotransmitter-Choline |
| Chrm4 | neurotransmitter-Choline |
| Chrm5 | neurotransmitter-Choline |
| Chrna1 | neurotransmitter-Choline |
| Chrna10 | neurotransmitter-Choline |
| Chrna2 | neurotransmitter-Choline |
| Chrna3 | neurotransmitter-Choline |
| Chrna4 | neurotransmitter-Choline |
| Chrna5 | neurotransmitter-Choline |
| Chrna6 | neurotransmitter-Choline |
| Chrna7 | neurotransmitter-Choline |
| Chrna9 | neurotransmitter-Choline |
| Chrnb1 | neurotransmitter-Choline |
| Chrnb2 | neurotransmitter-Choline |
| Chrnb3 | neurotransmitter-Choline |
| Chrnb4 | neurotransmitter-Choline |
| Chrnd | neurotransmitter-Choline |
| Chrne | neurotransmitter-Choline |
| Chrng | neurotransmitter-Choline |
| Cnksr3 | Oligo-Progen-2 |

| | |
|---|---|
| Cnr1 | neurotransmitter-anademide/cannabionoid |
| Cnr2 | neurotransmitter-anademide/cannabionoid |
| Col23a1 | Astrocytes-2 |
| Col25a1 | Meninges-2 |
| Col5a2 | DH Neurons |
| Cort | NP-GPCR/NPR genes |
| Cpne4 | DH Neurons |
| Crh | NP-GPCR/NPR genes |
| Crhr1 | NP-GPCR/NPR genes |
| Crhr2 | NP-GPCR/NPR genes |
| Ctss | Microglia |
| Cyp27a1 | Oligodendrocyte-2 |
| Dach2 | DH Neurons |
| Dbh | neurotransmitter-Norepinephrine and epinephrine |
| Dcbld2 | Ligand-receptor pairs-Semaphorin |
| Dcx | DH Neurons |
| Ddc | neurotransmitter-dopamine |
| Ddr1 | Ligand-receptor pairs-neurotrophin |
| Dlk1 | Ligand-receptor pairs-Differential-Dll3/Notch |
| Dlk2 | Ligand-receptor pairs-Differential-Dll3/Notch |
| Dll1 | Ligand-receptor pairs-Differential-Dll3/Notch |
| Dll3 | Ligand-receptor pairs-Differential-Dll3/Notch |
| Dll4 | Ligand-receptor pairs-Differential-Dll3/Notch |
| Dnah12 | Ependymal |
| Dpy19l1 | Oligodendrocyte-1 |
| Drd1 | neurotransmitter-dopamine |
| Drd2 | neurotransmitter-dopamine |
| Drd3 | neurotransmitter-dopamine |
| Drd4 | neurotransmitter-dopamine+Ligand-receptor pairs-neurotrophin |
| Drd5 | neurotransmitter-dopamine |
| Enpp1 | DH Neurons |
| Erbb2 | Ligand-receptor pairs-Semaphorin |
| Esrrb | VH Neuron |
| Esrrg | DH Neurons |
| Farp2 | Ligand-receptor pairs-Semaphorin |
| Fbn2 | MN |
| Fev | neurotransmitter-serotonine |
| Flt1 | Endothelial |
| Fos | Activity relavant genes |
| Fosb | Activity relavant genes |
| Foxp2 | VH Neuron |
| Fpr1 | Ligand-receptor pairs-TAFA4 |
| Gabbr2 | neurotransmitter-GABA |
| Gabra1 | DH Neurons+neurotransmitter-GABA |

| | |
|---|---|
| Gabra2 | DH Neurons+neurotransmitter-GABA |
| Gabra3 | neurotransmitter-GABA |
| Gabra4 | neurotransmitter-GABA |
| Gabra5 | neurotransmitter-GABA |
| Gabra6 | neurotransmitter-GABA |
| Gabrb1 | neurotransmitter-GABA |
| Gabrb2 | neurotransmitter-GABA |
| Gabrb3 | neurotransmitter-GABA |
| Gabrd | neurotransmitter-GABA |
| Gabre | neurotransmitter-GABA |
| Gabrg1 | neurotransmitter-GABA |
| Gabrg2 | neurotransmitter-GABA |
| Gabrg3 | neurotransmitter-GABA |
| Gabrp | neurotransmitter-GABA |
| Gabrq | neurotransmitter-GABA |
| Gabrr1 | neurotransmitter-GABA |
| Gabrr2 | neurotransmitter-GABA |
| Gabrr3 | neurotransmitter-GABA |
| Gad1 | neurotransmitter-GABA |
| Gad2 | neurotransmitter-GABA |
| Gal | DH Neurons |
| Galp | Neuroactive Peptides& Ligand-receptor pairs-Differential-Galanin |
| Galr1 | Neuroactive Peptides& Ligand-receptor pairs-Differential-Galanin |
| Galr2 | Neuroactive Peptides& Ligand-receptor pairs-Differential-Galanin |
| Galr3 | Neuroactive Peptides& Ligand-receptor pairs-Differential-Galanin |
| Gata3 | VH Neuron |
| Gbx2 | DH Neurons |
| Gch1 | neurotransmitter-serotonine |
| Gchfr | neurotransmitter-serotonine |
| Gfra3 | preganglionic cells (PGC) |
| Ghrl | Neuroactive Peptides-ghrelin |
| Ghsr | Neuroactive Peptides-ghrelin |
| Glra1 | neurotransmitter-glycine |
| Glra2 | neurotransmitter-glycine |
| Glra3 | neurotransmitter-glycine |
| Glra4 | neurotransmitter-glycine |
| Glrb | neurotransmitter-glycine |
| Gm26673 | DH Neurons |
| Gpc3 | DH Neurons |
| Gpr151 | Neuroactive Peptides& Ligand-receptor pairs-Differential-Galanin |
| Gpr35 | Neuroactive receptor-purinergic GPCR |
| Gpr83 | Neuroactive Peptides-Pen |
| Gria1 | neurotransmitter-glutamate |
| Gria3 | neurotransmitter-glutamate |

| | |
|---|---|
| Gria4 | neurotransmitter-glutamate |
| Grid1 | neurotransmitter-glutamate |
| Grid2 | neurotransmitter-glutamate |
| Grik1 | neurotransmitter-glutamate |
| Grik2 | neurotransmitter-glutamate |
| Grik3 | neurotransmitter-glutamate |
| Grik4 | neurotransmitter-glutamate |
| Grik5 | neurotransmitter-glutamate |
| Grin1 | neurotransmitter-glutamate |
| Grin2a | neurotransmitter-glutamate |
| Grin2b | neurotransmitter-glutamate |
| Grin2c | neurotransmitter-glutamate |
| Grin2d | neurotransmitter-glutamate |
| Grin3a | neurotransmitter-glutamate |
| Grin3b | neurotransmitter-glutamate |
| Grp | DH Neurons+NP-GPCR/NPR genes |
| Grpr | NP-GPCR/NPR genes |
| Hc | Neuroactive Peptides-Anaphylatoxin and TLQP-21 |
| Hcrt | Neuroactive Peptides-Orexin |
| Hcrtr1 | Neuroactive Peptides-Orexin |
| Hcrtr2 | Neuroactive Peptides-Orexin |
| Hdc | neurotransmitter-histamine |
| Hnmt | neurotransmitter-histamine |
| Hrh1 | neurotransmitter-histamine |
| Hrh2 | neurotransmitter-histamine |
| Hrh3 | neurotransmitter-histamine |
| Hrh4 | neurotransmitter-histamine |
| Htr1a | neurotransmitter-serotonine |
| Htr1b | neurotransmitter-serotonine |
| Htr1d | neurotransmitter-serotonine |
| Htr1f | neurotransmitter-serotonine |
| Htr2a | neurotransmitter-serotonine |
| Htr2b | neurotransmitter-serotonine |
| Htr2c | neurotransmitter-serotonine |
| Htr3a | neurotransmitter-serotonine |
| Htr3b | neurotransmitter-serotonine |
| Htr4 | neurotransmitter-serotonine |
| Htr5a | neurotransmitter-serotonine |
| Htr5b | neurotransmitter-serotonine |
| Htr6 | neurotransmitter-serotonine |
| Htr7 | neurotransmitter-serotonine |
| Iapp | Ligand-receptor pairs-CGRP |
| Ier2 | cell growth |
| Itga1 | Ligand-receptor pairs-Semaphorin |

| | |
|---|---|
| Itgb1 | Ligand-receptor pairs-Semaphorin |
| Jag1 | Ligand-receptor pairs-Differential-Dll3/Notch |
| Jag2 | Ligand-receptor pairs-Differential-Dll3/Notch |
| Junb | Activity relavant genes |
| Kcnip2 | DH Neurons |
| Kidins220 | Ligand-receptor pairs-neurotrophin |
| Klhl14 | DH Neurons |
| Kng1 | Neuroactive Peptides-Bradykinin |
| Kng2 | Neuroactive Peptides-Bradykinin |
| Lhx2 | DH Neurons |
| Lhx9 | DH Neurons |
| Lmx1b | DH Neurons |
| Lpar4 | Neuroactive receptor-purinergic GPCR |
| Lpar6 | Neuroactive receptor-purinergic GPCR |
| Ly6c1 | Endothelial |
| Maf | DH Neurons |
| Mc1r | Neuroactive Peptides-Melanocortin |
| Mc2r | Neuroactive Peptides-Melanocortin |
| Mc3r | Neuroactive Peptides-Melanocortin |
| Mc4r | Neuroactive Peptides-Melanocortin |
| Mc5r | Neuroactive Peptides-Melanocortin |
| Mdga1 | DH Neurons |
| Megf11 | DH Neurons |
| Met | Ligand-receptor pairs-Semaphorin |
| Mlxipl | DH Neurons |
| Myo1f | Microglia |
| Neurod1 | generic neuronal marker |
| Neurod2 | DH Neurons |
| Nfib | DH Neurons |
| Ngf | Ligand-receptor pairs-neurotrophin |
| Ngfr | Ligand-receptor pairs-neurotrophin |
| Nmb | NP-GPCR/NPR genes |
| Nmbr | NP-GPCR/NPR genes |
| Nms | DH Neurons |
| Nmu | DH Neurons |
| Nmur1 | Neuroactive Peptides-Neuromedin U/S |
| Nmur2 | DH Neurons+Neuroactive Peptides-Neuromedin U/S |
| Nos1 | preganglionic cells (PGC)+DH Neurons |
| Notch1 | Ligand-receptor pairs-Differential-Dll3/Notch |
| Notch2 | Ligand-receptor pairs-Differential-Dll3/Notch |
| Notch3 | Pericytes+Ligand-receptor pairs-Differential-Dll3/Notch |
| Notch4 | Ligand-receptor pairs-Differential-Dll3/Notch |
| Npas4 | Activity relavant genes |
| Npb | Neuroactive Peptides-Neuropeptide W/B |

| | |
|---|---|
| Npbwr1 | Neuroactive Peptides-Neuropeptide W/B |
| Npff | Neuroactive Peptides-Neuropeptide FF/VF |
| Npffr1 | Neuroactive Peptides-Neuropeptide FF/VF |
| Npffr2 | Neuroactive Peptides-Neuropeptide FF/VF |
| Nppc | DH Neurons |
| Nps | Neuroactive Peptides-Neuropeptide S |
| Npsr1 | Neuroactive Peptides-Neuropeptide S |
| Npvf | Neuroactive Peptides-Neuropeptide FF/VF |
| Npw | Neuroactive Peptides-Neuropeptide W/B |
| Npy | DH Neurons+NP-GPCR/NPR genes |
| Npy1r | NP-GPCR/NPR genes |
| Npy2r | NP-GPCR/NPR genes |
| Npy4r | NP-GPCR/NPR genes |
| Npy5r | NP-GPCR/NPR genes |
| Nr4a1 | Activity relavant genes |
| Nr5a2 | VH Neuron |
| Nrgn | DH Neurons |
| Nrp1 | Ligand-receptor pairs-Semaphorin |
| Nrp2 | Ligand-receptor pairs-Semaphorin |
| Ntf3 | Ligand-receptor pairs-neurotrophin |
| Ntf5 | Ligand-receptor pairs-neurotrophin |
| Ntrk1 | Ligand-receptor pairs-neurotrophin |
| Ntrk2 | Ligand-receptor pairs-neurotrophin |
| Ntrk3 | Ligand-receptor pairs-neurotrophin |
| Nts | DH Neurons |
| Ntsr1 | NP-GPCR/NPR genes |
| Ntsr2 | NP-GPCR/NPR genes |
| Onecut2 | DH Neurons |
| Opalin | Oligo-Progen-2 |
| Oprd1 | NP-GPCR/NPR genes |
| Oprk1 | NP-GPCR/NPR genes |
| Oprl1 | NP-GPCR/NPR genes |
| Oprm1 | NP-GPCR/NPR genes |
| Oxtr | Neuroactive Peptides-Oxytocin |
| P2rx1 | Neuroactive receptor-ATP Receptor/purinoceptor |
| P2rx2 | Neuroactive receptor-ATP Receptor/purinoceptor |
| P2rx3 | Neuroactive receptor-ATP Receptor/purinoceptor |
| P2rx4 | Neuroactive receptor-ATP Receptor/purinoceptor |
| P2rx5 | Neuroactive receptor-ATP Receptor/purinoceptor |
| P2rx6 | Neuroactive receptor-ATP Receptor/purinoceptor |
| P2rx7 | Neuroactive receptor-ATP Receptor/purinoceptor |
| P2ry1 | Neuroactive receptor-purinergic GPCR |
| P2ry10 | Neuroactive receptor-purinergic GPCR |
| P2ry10b | Neuroactive receptor-purinergic GPCR |

| | |
|---|---|
| P2ry13 | Neuroactive receptor-purinergic GPCR |
| P2ry14 | Neuroactive receptor-purinergic GPCR |
| P2ry2 | Neuroactive receptor-purinergic GPCR |
| P2ry4 | Neuroactive receptor-purinergic GPCR |
| P2ry6 | Neuroactive receptor-purinergic GPCR |
| Pax2 | VH Neuron |
| Pax6 | DH Neurons |
| Pcdha1 | clustered Pcdhs |
| Pcdha11 | clustered Pcdhs |
| Pcdha12 | clustered Pcdhs |
| Pcdha2 | clustered Pcdhs |
| Pcdha3 | clustered Pcdhs |
| Pcdha4 | clustered Pcdhs |
| Pcdha5 | clustered Pcdhs |
| Pcdha6 | clustered Pcdhs |
| Pcdha7 | clustered Pcdhs |
| Pcdha8 | clustered Pcdhs |
| Pcdha9 | clustered Pcdhs |
| Pcdhac1 | clustered Pcdhs |
| Pcdhac2 | clustered Pcdhs |
| Pcdhb1 | clustered Pcdhs |
| Pcdhb10 | clustered Pcdhs |
| Pcdhb11 | clustered Pcdhs |
| Pcdhb12 | clustered Pcdhs |
| Pcdhb13 | clustered Pcdhs |
| Pcdhb14 | clustered Pcdhs |
| Pcdhb15 | clustered Pcdhs |
| Pcdhb16 | clustered Pcdhs |
| Pcdhb17 | clustered Pcdhs |
| Pcdhb18 | clustered Pcdhs |
| Pcdhb19 | clustered Pcdhs |
| Pcdhb2 | clustered Pcdhs |
| Pcdhb20 | clustered Pcdhs |
| Pcdhb21 | clustered Pcdhs |
| Pcdhb22 | clustered Pcdhs |
| Pcdhb3 | clustered Pcdhs |
| Pcdhb4 | clustered Pcdhs |
| Pcdhb5 | clustered Pcdhs |
| Pcdhb6 | clustered Pcdhs |
| Pcdhb7 | clustered Pcdhs |
| Pcdhb8 | clustered Pcdhs |
| Pcdhb9 | clustered Pcdhs |
| Pcdhga1 | clustered Pcdhs |
| Pcdhga10 | clustered Pcdhs |

| | |
|---|---|
| Pcdhga11 | clustered Pcdhs |
| Pcdhga12 | clustered Pcdhs |
| Pcdhga2 | clustered Pcdhs |
| Pcdhga3 | clustered Pcdhs |
| Pcdhga4 | clustered Pcdhs |
| Pcdhga5 | clustered Pcdhs |
| Pcdhga6 | clustered Pcdhs |
| Pcdhga7 | clustered Pcdhs |
| Pcdhga8 | clustered Pcdhs |
| Pcdhga9 | clustered Pcdhs |
| Pcdhgb1 | clustered Pcdhs |
| Pcdhgb2 | clustered Pcdhs |
| Pcdhgb4 | clustered Pcdhs |
| Pcdhgb5 | clustered Pcdhs |
| Pcdhgb6 | clustered Pcdhs |
| Pcdhgb7 | clustered Pcdhs |
| Pcdhgb8 | clustered Pcdhs |
| Pcdhgc3 | clustered Pcdhs |
| Pcdhgc4 | clustered Pcdhs |
| Pcdhgc5 | clustered Pcdhs |
| Pcsk1n | Neuroactive Peptides-Pen |
| Pdgfra | OPC |
| Pdyn | DH Neurons+NP-GPCR/NPR genes |
| Pdzrn3 | Meninges-1 |
| Penk | NP-GPCR/NPR genes |
| Pitx2 | DH Neurons |
| Pkd2l1 | DH Neurons |
| Plxna1 | Ligand-receptor pairs-Semaphorin |
| Plxna2 | Ligand-receptor pairs-Semaphorin |
| Plxna3 | Ligand-receptor pairs-Semaphorin |
| Plxna4 | Ligand-receptor pairs-Semaphorin |
| Plxnb1 | Ligand-receptor pairs-Semaphorin |
| Plxnb2 | Ligand-receptor pairs-Semaphorin |
| Plxnb3 | Ligand-receptor pairs-Semaphorin |
| Plxnc1 | Ligand-receptor pairs-Semaphorin |
| Plxnd1 | Ligand-receptor pairs-Semaphorin |
| Pmfbp1 | DH Neurons |
| Pmp22 | Schwann |
| Pnmt | neurotransmitter-Norepinephrine and epinephrine |
| Pnoc | NP-GPCR/NPR genes |
| Poln | MN |
| Pomc | Neuroactive Peptides-Melanocortin |
| Pou6f2 | DH Neurons |
| Prkcg | DH Neurons |

| | |
|---|---|
| Prlr | DH Neurons |
| Prox1 | DH Neurons |
| Ptgds | Oligodendrocyte-1 |
| Pth1r | NP-GPCR/NPR genes |
| Pthlh | NP-GPCR/NPR genes |
| Pts | neurotransmitter-serotonine |
| Pvalb | DH Neurons |
| Qdpr | neurotransmitter-serotonine |
| Qrfp | Neuroactive Peptides-Orexigenic Neuropeptide |
| Qrfpr | DH Neurons+Neuroactive Peptides-Orexigenic Neuropeptide |
| Rab37 | Oligodendrocyte-2 |
| Ramp1 | Ligand-receptor pairs-CGRP |
| Ramp2 | Ligand-receptor pairs-CGRP |
| Ramp3 | Ligand-receptor pairs-CGRP |
| Rbfox3 | generic neuronal marker |
| Reln | DH Neurons |
| Rln1 | NP-GPCR/NPR genes |
| Rnf220 | VH Neuron |
| Rorb | DH Neurons |
| Rreb1 | DH Neurons |
| Rspo3 | DH Neurons |
| Runx2 | DH Neurons |
| Rxfp1 | NP-GPCR/NPR genes |
| Rxfp2 | DH Neurons+NP-GPCR/NPR genes |
| Rxfp3 | NP-GPCR/NPR genes |
| Satb1 | DH Neurons |
| Satb2 | DH Neurons |
| Scn7a | Peripheral glia |
| Sema3a | Ligand-receptor pairs-Semaphorin |
| Sema3b | Ligand-receptor pairs-Semaphorin |
| Sema3c | Ligand-receptor pairs-Semaphorin |
| Sema3d | Ligand-receptor pairs-Semaphorin |
| Sema3e | Ligand-receptor pairs-Semaphorin |
| Sema3f | Ligand-receptor pairs-Semaphorin |
| Sema3g | Ligand-receptor pairs-Semaphorin |
| Sema4a | Ligand-receptor pairs-Semaphorin |
| Sema4b | Ligand-receptor pairs-Semaphorin |
| Sema4c | Ligand-receptor pairs-Semaphorin |
| Sema4d | Ligand-receptor pairs-Semaphorin |
| Sema4f | Ligand-receptor pairs-Semaphorin |
| Sema4g | Ligand-receptor pairs-Semaphorin |
| Sema5a | Ligand-receptor pairs-Semaphorin |
| Sema5b | DH Neurons+Ligand-receptor pairs-Semaphorin |
| Sema6a | Ligand-receptor pairs-Semaphorin |

| | |
|---|---|
| Sema6b | Ligand-receptor pairs-Semaphorin |
| Sema6c | Ligand-receptor pairs-Semaphorin |
| Sema6d | Ligand-receptor pairs-Semaphorin |
| Sema7a | Ligand-receptor pairs-Semaphorin |
| Shox2 | VH Neuron |
| Sim1 | VH Neuron |
| Slc17a6 | neurotransmitter-glutamate |
| Slc17a7 | neurotransmitter-glutamate |
| Slc17a8 | DH Neurons |
| Slc18a2 | neurotransmitter-serotonine |
| Slc18a3 | neurotransmitter-Choline |
| Slc32a1 | neurotransmitter-glycine |
| Slc47a1 | Meninges-2 |
| Slc6a1 | GABA transporter |
| Slc6a2 | neurotransmitter-dopamine |
| Slc6a3 | neurotransmitter-dopamine |
| Slc6a4 | neurotransmitter-serotonine |
| Slc6a5 | DH Neurons+neurotransmitter-glycine |
| Slc7a10 | Astrocytes-1 |
| Sorcs3 | DH Neurons+Ligand-receptor pairs-neurotrophin |
| Sort1 | Ligand-receptor pairs-neurotrophin |
| Sox5 | DH Neurons |
| Spp1 | MN |
| Spr | neurotransmitter-serotonine |
| Spx | Neuroactive Peptides& Ligand-receptor pairs-Differential-Galanin |
| Sst | NP-GPCR/NPR genes |
| Sstr1 | NP-GPCR/NPR genes |
| Sstr2 | NP-GPCR/NPR genes |
| Sstr3 | NP-GPCR/NPR genes |
| Sstr4 | NP-GPCR/NPR genes |
| Syt2 | DH Neurons |
| Tac1 | DH Neurons+NP-GPCR/NPR genes |
| Tac2 | DH Neurons |
| Tac4 | NP-GPCR/NPR genes |
| Tacr1 | NP-GPCR/NPR genes |
| Tacr2 | NP-GPCR/NPR genes |
| Tacr3 | NP-GPCR/NPR genes |
| Tafa4 | Ligand-receptor pairs-TAFA4 |
| Tfap2b | DH Neurons |
| Th | neurotransmitter-dopamine |
| Tnr | Oligo-Progen-1 |
| Tns1 | MN |
| Tph1 | neurotransmitter-serotonine |
| Tph2 | neurotransmitter-serotonine |

| | |
|---|---|
| Trem2 | Ligand-receptor pairs-Semaphorin |
| Trh | NP-GPCR/NPR genes |
| Trhr | DH Neurons |
| Trhr2 | NP-GPCR/NPR genes |
| Trp63 | NP-GPCR/NPR genes |
| Tyrobp | Ligand-receptor pairs-Semaphorin |
| Ust | Oligo-Progen-1 |
| Uts2 | Neuroactive Peptides-Urotensin II |
| Uts2b | Neuroactive Peptides-Urotensin II |
| Uts2r | Neuroactive Peptides-Urotensin II |
| Vamp1 | DH Neurons |
| Vgf | DH Neurons+Neuroactive Peptides-Anaphylatoxin and TLQP-21 |
| Vip | NP-GPCR/NPR genes |
| Vipr1 | NP-GPCR/NPR genes |
| Vipr2 | NP-GPCR/NPR genes |
| Vsx2 | DH Neurons |
| Zfhx3 | DH Neurons |
| Zim1 | DH Neurons |

Table S2 Proportions of neuronal subtypes in the spinal cord.

| Neuron subtype | Cervical | Thoracic | Lumbar | Sacral |
|---|---|---|---|---|
| DM-ex-Zfhx3 | 0.0661 | 0.0661 | 0.0623 | 0.0652 |
| MV-ex-Syt2 | 0.0534 | 0.0433 | 0.0658 | 0.0398 |
| DH-ex-Reln/Nmur2 | 0.0341 | 0.0406 | 0.0300 | 0.0289 |
| DH-ex-Nmu/Tac2 | 0.0277 | 0.0378 | 0.0199 | 0.0184 |
| DH-ex-Gpr83 | 0.0230 | 0.0138 | 0.0184 | 0.0110 |
| DH-ex-Maf/Cck | 0.0264 | 0.0254 | 0.0226 | 0.0270 |
| DH-ex-Prkcg/Nts | 0.0289 | 0.0273 | 0.0184 | 0.0125 |
| DH-ex-Prkcg/Cck | 0.0219 | 0.0165 | 0.0178 | 0.0219 |
| DH-ex-Reln | 0.0173 | 0.0149 | 0.0159 | 0.0136 |
| MV-ex-Shox2 | 0.0205 | 0.0214 | 0.0194 | 0.0192 |
| DH-ex-Reln/Npff | 0.0144 | 0.0156 | 0.0108 | 0.0124 |
| DH-ex-Sox5 | 0.0192 | 0.0288 | 0.0199 | 0.0222 |
| DH-ex-Tac2 | 0.0169 | 0.0176 | 0.0175 | 0.0186 |
| DH-ex-Sox5/Tac1 | 0.0113 | 0.0153 | 0.0100 | 0.0155 |
| DH-ex-Cpne4 | 0.0161 | 0.0126 | 0.0114 | 0.0121 |
| DH-ex-Rreb1 | 0.0099 | 0.0092 | 0.0089 | 0.0112 |
| DH-ex-Prkcg/Rxfp1 | 0.0142 | 0.0194 | 0.0191 | 0.0132 |
| DH-ex-Grp | 0.0115 | 0.0101 | 0.0103 | 0.0141 |
| M-ex-Vsx2/Shox2 | 0.0151 | 0.0113 | 0.0188 | 0.0149 |
| M-ex-Vsx2 | 0.0168 | 0.0150 | 0.0245 | 0.0274 |
| DH-ex-Grpr | 0.0140 | 0.0156 | 0.0105 | 0.0100 |
| DH-ex-Maf/Cpne4 | 0.0044 | 0.0054 | 0.0035 | 0.0059 |
| M-ex-Neurod2 | 0.0019 | 0.0093 | 0.0064 | NaN |
| DH-ex-Maf/Slc17a8 | 0.0055 | 0.0084 | 0.0057 | 0.0013 |
| MV-in-Gabra1 | 0.0667 | 0.0662 | 0.0685 | 0.0734 |
| DH-in-Cdh3 | 0.0599 | 0.0549 | 0.0534 | 0.0679 |
| MV-in-Gm26673 | 0.0526 | 0.0425 | 0.0634 | 0.0479 |
| M-in-Tfap2b | 0.0283 | 0.0350 | 0.0347 | 0.0243 |
| MV-in-Esrrb | 0.0321 | 0.0279 | 0.0378 | 0.0582 |
| DH-in-Npy | 0.0238 | 0.0250 | 0.0175 | 0.0246 |
| DH-in-Pdyn/Gal | 0.0155 | 0.0172 | 0.0109 | 0.0132 |
| MV-in-Sema5b | 0.0478 | 0.0378 | 0.0497 | 0.0898 |
| DH-in-Npy2r | 0.0141 | 0.0135 | 0.0159 | 0.0224 |
| DH-in-Pdyn | 0.0244 | 0.0260 | 0.0218 | 0.0150 |
| DH-in-Rorb | 0.0162 | 0.0200 | 0.0147 | 0.0097 |
| MV-in-Chrna2 | 0.0163 | 0.0189 | 0.0185 | 0.0266 |
| DH-in-Kcnip2 | 0.0066 | 0.0064 | 0.0067 | 0.0091 |
| DH-in-Klhl14 | 0.0095 | 0.0080 | 0.0101 | 0.0083 |
| M-in-Chat | 0.0016 | 0.0012 | 0.0016 | NaN |
| alpha motoneuron | 0.0611 | 0.0474 | 0.0602 | 0.0386 |
| beta motoneuron | 0.0118 | 0.0122 | 0.0158 | 0.0072 |
| gamma motoneuron | 0.0139 | 0.0179 | 0.0224 | 0.0272 |

| | | | | |
|---|---|---|---|---|
| cholinergic interneuron | 0.0029 | 0.0040 | 0.0019 | NaN |
| visceral motoneuron | 0.0043 | 0.0174 | 0.0067 | 0.0002 |

**Supplemental Information References**
